# A large European diversity panel reveals complex azole fungicide resistance gains of a major wheat pathogen

**DOI:** 10.1101/2025.03.19.644200

**Authors:** Guido Puccetti, Thomas Badet, Daniel Flückiger, Dominique Edel, Alice Feurtey, Camille Delude, Emile-Gluck Thaler, Stefano F.F. Torriani, Gabriel Scalliet, Daniel Croll

## Abstract

Fungicide resistance in crop pathogens poses severe challenges to sustainable agriculture. Demethylation inhibitors (DMIs) are critical for controlling crop diseases but face rapid resistance gains in the field. Even though the main molecular basis of resistance is well established, field surveys have repeatedly revealed alternative resistance mechanisms. The European continent in particular has seen rapid and heterogeneous gains in azole resistance in the past decades. Here, we establish a large genome panel to dissect the genetic architecture of emerging resistance in the major wheat pathogen *Zymoseptoria tritici*. The European diversity panel spans 15 sampling years and 27 countries for a total of 1394 sequenced and phenotyped strains. Using two complementary assays to quantify resistance levels of each strain, we captured fine-grained shifts in DMI resistance over space and time. We conducted genome-wide association studies based on a comprehensive set of genotyping approaches for six DMIs. We mapped a total of 21,220 genetic variants and 158 genes linked to resistance. The substantial scope in genetic mechanisms underpinning DMI resistance significantly expands our mechanistic understanding how continent-wide resistance arises in fungal pathogens over time. Diversification of the *Cyp51* coding sequence was particularly striking with new resistant haplotypes emerging with complex configurations and geographic patterns. This study provides expansive new insights into fungicide resistance gains of crop pathogens relevant for future resistance management strategies.

## Introduction

The application of fungicides in agriculture is the primary containment strategy to reduce crop losses caused by fungal pathogens (1). However, resistance can rise rapidly in fields following intense application of fungicides (2). The speed of resistance gains and the geographic scope remain difficult to predict (2, 3). The emergence of fungicide resistance highlights the challenges of convergent adaptation across geographic space (4). Even though the molecular basis of fungicide resistance may be well established for specific pathogens and fungicides, field surveys have repeatedly revealed alternative resistance mechanisms (5–7). Demethylation inhibitors (DMIs) are among the most widely applied fungicides (8, 9). Resistance to this class of fungicides is mainly attributed to mutations in the gene encoding the molecular target *Cyp51*, which encodes the enzyme sterol 14α-demethylase. This enzyme is essential for the ergosterol biosynthesis pathway underpinning the production of sterols for fungal cell membranes (10, 11). A number of recent studies showed that DMI resistance is also mediated by additional mechanisms beyond mutations in the *Cyp51* gene such as regulatory changes in promoter elements leading to overexpression (12), as well as non-target resistance caused by the overexpression of efflux pumps (13, 14). In crop pathogens, field resistance to DMIs has risen due to a combination of different on and off-target mechanisms (6, 15). Off-target mechanisms, such as the combinatorial effects of multiple genetic factors, have been particularly significant. For instance, resistance to fluconazole in *Candida albicans* can be gained in complex steps mediated by overexpressed NCP1, cytochrome p450, and loss of heterozygosity at *KSR1* involved in sphingolipid biosynthesis in addition to chromosome copy-number variation (16). However, it remains largely unknown how such combinatorial effects of resistance mutations contribute to the gain of fungicide resistance over space and time in the field.

Population genomic surveys of pathogen populations have facilitated tracking fungicide resistance against DMIs and other fungicides at the field and continental scales. Recent gains in DMI resistance in the fungal wheat pathogen *Zymoseptoria tritici* were likely underpinned by both convergent and unique resistance adaptations (17, 18). Resistance to the DMI propiconazole was associated with mutations in a gene encoding a DHHC palmitoyl transferase and were geographically restricted to fields in the United States (17). Geographic variation in fungicide resistance has also been observed in *Fusarium fujikuroi* with differing sensitivity levels reported between countries (19, 20). Regional differences in the rise of resistance towards succinate dehydrogenase inhibitor (SDHI) fungicides were likely in part caused by variation in selection strength and geographic isolation (21–24). Key insights into population-level resistance mutations were gained using genome-wide association studies (GWAS) associating phenotypic trait variation (*i.e.* fungicide resistance) to molecular variation (25–28). Applications of GWAS in *Z. tritici*, the barley scald pathogen *Rhynchosporium commune* and *Cercospora beticola* causing Cercospora leaf spot revealed both major contributions of *Cyp51* mutations, but also previously unknown loci contributing to DMI resistance (17, 27–29). GWAS further revealed complex associations between *Cyp51* mutations and levels of resistance towards different DMIs fungicides. In *C. beticola,* a synonymous single nucleotide polymorphism (SNP) was associated with tetraconazole resistance as well as a further synonymous SNP underpinning *Cyp51* expression variation (28). However, significant associations of synonymous variants could also be explained by high linkage disequilibrium (LD), especially following a selective sweep for resistance gains.

Assessing fungicide resistance experimentally is highly sensitive to the composition of the test medium, temperature, and availability of the fungicide in the medium. Resistance is commonly assessed in liquid culture medium over an array of fungicide doses (30–32). However, many filamentous fungi do not grow well in submerged cultures as they show unusual morphologies including dispersed hyphae or form pellets (33, 34). In contrast, solid media assays typically require larger amounts of fungicides compared to liquid-based methods and are challenging to standardize (35, 36). Further complicating fungicide resistance phenotyping efforts are inconsistencies in resistance assessed in liquid versus solid media. Such issues were detected *e.g.* for resistance assessments against propiconazole in *Clarireedia jacksonii* (37). Hence, high-throughput assessment of among individual variation in resistance requires efforts in standardization and protocol development.

With the widespread application of DMIs on the European continent, fungal plant pathogens such as the wheat pathogen *Z. tritici* appeared as major threats given their ability to evolve resistance (38, 39). European wheat production heavily relies on DMIs for control to prevent yield losses on the order of 50% (40). *Z. tritici* populations show remarkable evolutionary potential to overcome host resistance and application of fungicides. Adaptation is facilitated by high levels of genetic variation at the field and continental scale (26, 41), including structural variation and accessory chromosomes (42–44). Furthermore, high rates of recombination (45, 46) and the activity of transposable elements (TEs) contribute to highly diverse populations (47, 48). Gains in DMI resistance was in part mediated by numerous point mutations in the target gene *Cyp51* (49–53). *Z. tritici* populations are characterized by a complex mixture of resistant *Cyp51* haplotypes underlying a differential effect of mutations across DMIs (50). Some key non-synonymous mutations include Ser524Thr, which increases resistance against at least four DMIs (50, 53, 54). The Ile381Val substitution underpins resistance against two additional DMIs (55). *Z. tritici* evolved also non-target site mechanisms to tolerate DMIs (6, 17, 56, 57). The complexity in mutations contributing to DMI resistance raises significant questions about the geographic scope and convergence in resistance gains. Such lack of knowledge limits also our ability to predict the emergence of resistance and protect crops effectively.

Here, we established and investigated the largest hierarchically collected sampling of *Z. tritici* across the European continent covering a time span of 15 years. We optimized phenotyping assays to comprehensively capture shifts in DMI fungicide resistance across 1394 strains. GWAS conducted for 6 DMIs identified 21,220 mutations across 158 genes contributing to resistance among DMIs. Resistance mutations were gained heterogeneously among the major wheat producing areas. Finally, we assessed contributions of coding sequence mutations in *Cyp51* and validated a resistance shift caused by a specific emerging *Cyp51* haplotype.

## Results

### A European diversity panel to survey resistance emergence

To investigate the genetic basis of DMI resistance under field conditions, we selected strains collected on infected wheat leaves across diverse regions in Europe. Sampling was particularly dense in Central and Northern European regions of intense wheat production and exposure to more fungicide applications (58). strains originated from 27 countries sampled over a time span of 15 years (Figure 1A) (Table S1). Strains from Germany, France, the U.K., and Ireland accounted for ∼72% of the panel (Figure 1B). The retained diversity panel was following a hierarchical sampling approach favoring strains from distinct wheat fields, at larger geographic distances and spread over the available sampling years (see Methods). This selection process maximized the covered geographic area and timespan.

**Figure 1.**
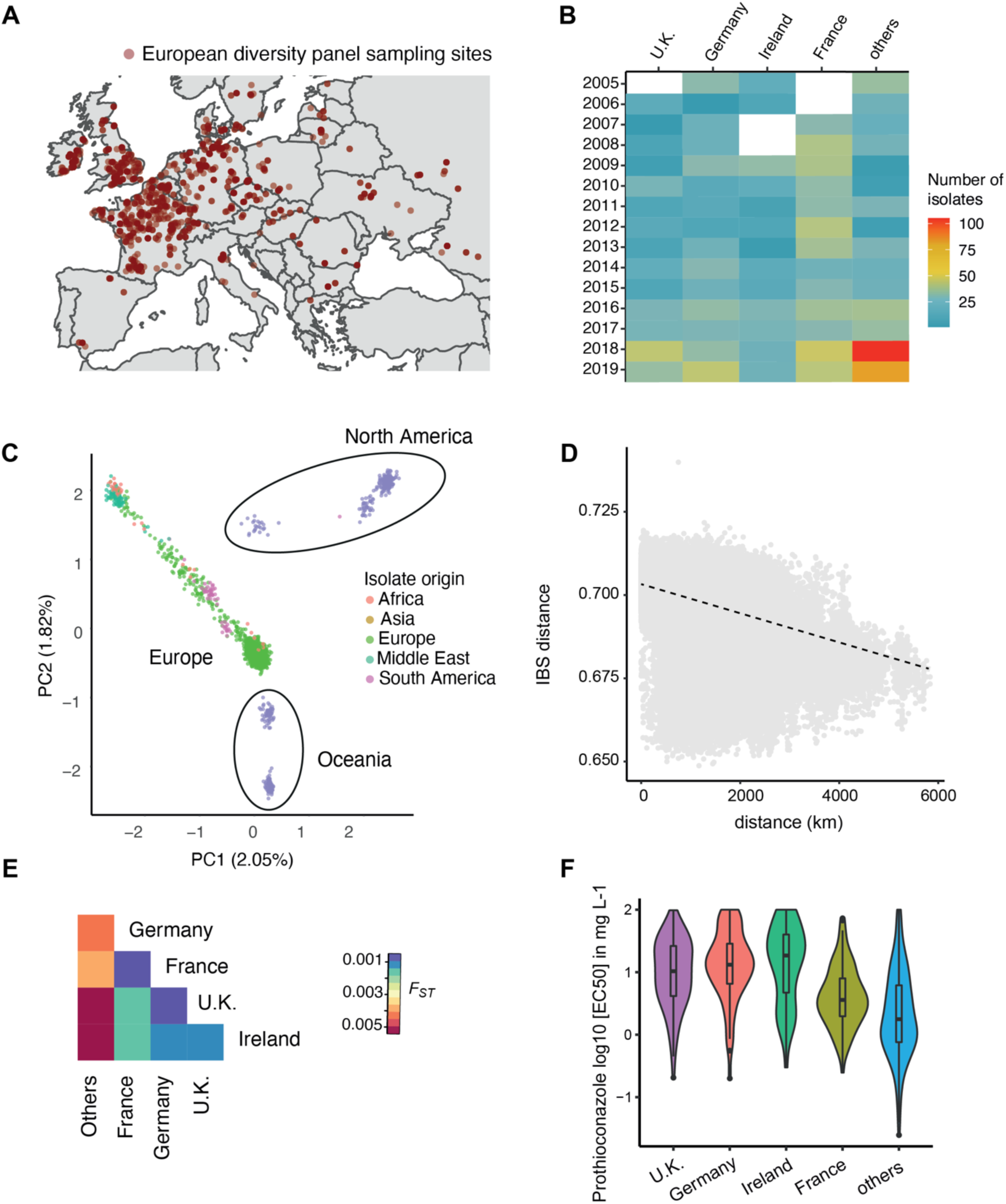
Collection and analyses of the fungal wheat pathogen *Zymoseptoria tritici* to create a European diversity panel (*n* = 1394 strains). (A) Sampling distribution across Europe of the European diversity panel. (B) Heatmap representing the number of strains sampled in France, Germany, U.K., Ireland and the rest of Europe. (C) Principal component analysis of 2156 strains including a separate strain collection covering additional continents. (D) Isolation-by-distance analyses using identity-by-state (IBS) pairwise distances. (E) Heatmap of the FST values for strains sampled from Germany, France, U.K., Ireland, and Others. (F) EC50 values of prothioconazole for strains sampled in Germany, U.K., France, Ireland, and the rest of Europe in the years 2016-2018.

Each *Z. tritici* strain of the European diversity panel was prepared for long-term preservation, phenotypic assays and whole-genome Illumina sequencing. We followed a previously established genotyping and filtering procedure to obtain the most robust set of SNP and short indel variants (26). To assess the genetic diversity of the European panel in the global genetic context, we analyzed the newly generated data jointly with existing genotyping data from a thousand-genome panel of the species (26). This resulted in a combination of 1,134 newly and 1,022 previously sequenced datasets for a total of 2,156 high-quality genomes. We identified 8,536,499 raw SNP and 6,606,877 raw indel candidate loci. After stringent quality filtering, we retained 472,041 SNPs for the European diversity panel with minor allele frequencies of at least 5% for further processing. The global genetic diversity of the species was primarily structured into a North American, an Oceanian and a dispersed third group including strains originating from Europe, the Middle East, North Africa and South America, consistent with previous genetic diversity analyses (26) (Figure 1C). The European diversity panel showed a striking spread in genetic diversity consistent with Europe being a founder population for other continents as well as experiencing recent incoming gene flow.

We analyzed fine-scale admixture patterns within the European diversity panel and identified 11 clusters with a broad geographic spread (Figure S1B-D). Gene flow has occurred over large distances as indicated by a slow decay of identity-by-state (IBS) over geographic distance (Figure 1D). Genetic differentiation among major wheat producers, *i.e.* Germany, France, the U.K., Ireland was generally low consistent with extensive gene flow across Europe (Figure 1E). To assess mapping power in the panel, we investigated LD decay and found that *r*² decayed to ∼0.2 within ∼400 bp (Figure S2B). Rapid LD decay is a hallmark of highly diverse populations with high effective population size.

### Profiling fungicide resistance across the continent

We assessed resistance in the panel against DMIs including prothioconazole, prochloraz, epoxiconazole, tebuconazole, and cyproconazole, which are all applied in Europe to control *Z. tritici* (Table S2) (European Commission, 2024). Half maximal effective concentration (EC_50_) values were notably higher in strains from Western and Northern Europe (*i.e.*, France, Ireland, Germany, and the U.K.) compared to the rest of the continent (Figure 1F). From 2005-2019, resistance increased across most sampled regions consistent with broad fungicide resistance gains on the continent (Figure S1A). We estimated fungicide resistance using growth assessments in two media: relative colony size on solid media with or without fungicide in the medium (Figure 2A) and by determining half inhibitory concentrations (EC_50_) inferred from liquid cultures at an array of concentrations (Figure 2B). Resistance assessed using the two assays showed correlation *r* = 0.61-0.67 except for prochloraz (*r* = 0.37; Figure S3A-D). The largest discrepancies were observed for some strains with high EC_50_ values (>10 mg.L^-1^), which did not exhibit growth on solid media. For instance, in cyproconazole 306 strains showed no growth on solid medium at a concentration of 10 mg.L^-1^, yet their EC₅₀ values in liquid assays ranged from 0.018-71.05 mg.L-1. Similarly a wide spectrum of fungicide resistance (0.007-72.0 mg.L-1) was observed for prothioconazole in liquid assays and 311 strains showed no growth on solid medium at a concentration of 1 mg.L-1. Concordance among DMI fungicides was high (*i.e.* cross-resistance) with discrepancies restricted to specific fungicide pairs including prothioconazole and tebuconazole, prothioconazole and prochloraz, and tebuconazole and prochloraz (Figure 2C-D). These pairs showed positively correlated resistance levels as assessed by EC_50_ values. The same pairs showed only weak or no resistance correlation on solid medium. The divergence highlights how resistance assays can influence the detection of cross-resistance patterns. The observed discrepancies highlight the relevance of assessing fungicide resistance in multiple morphological states spanning a range of concentrations.

**Figure 2.**
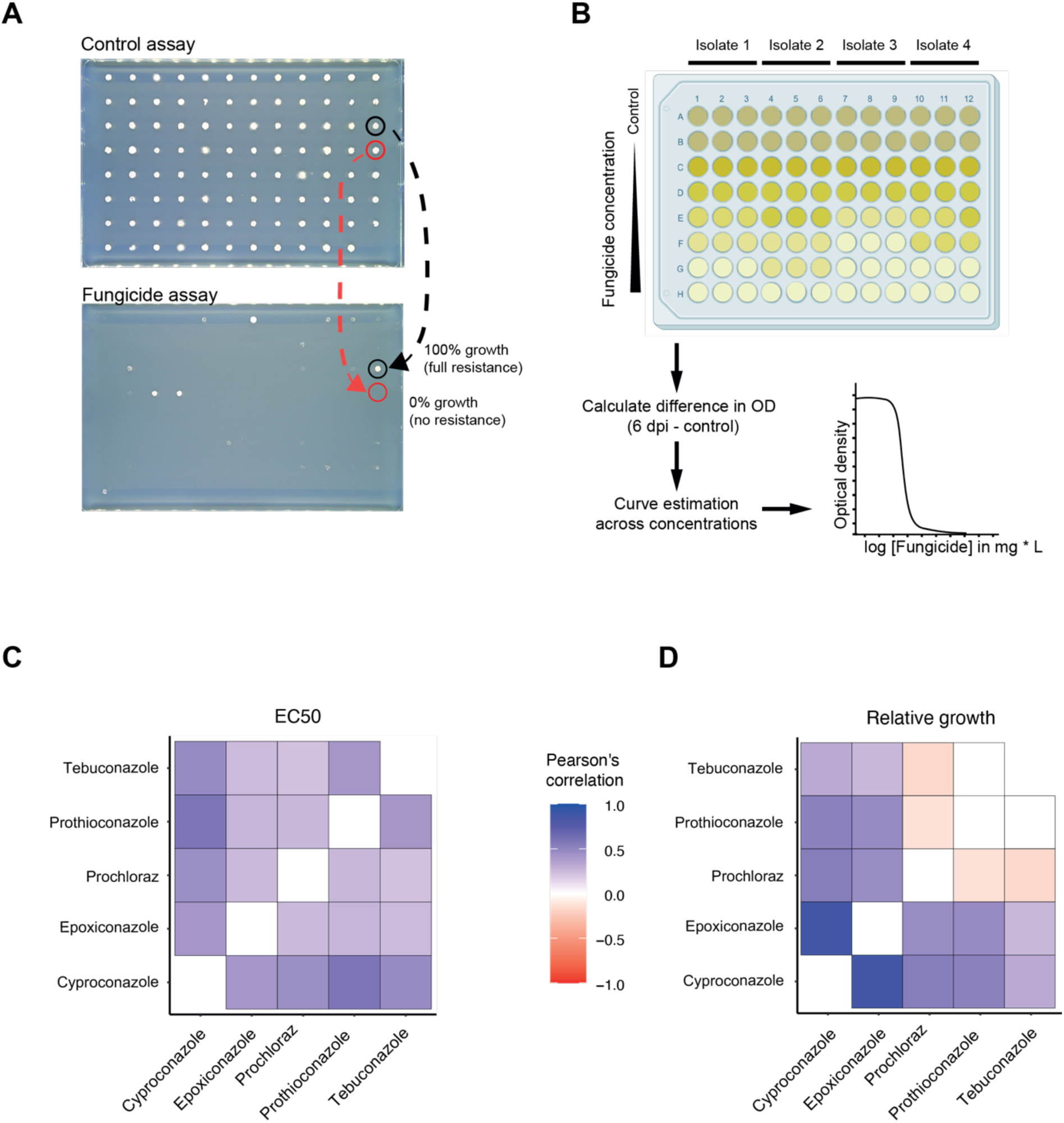
Phenotyping assays and cross-resistance among demethylation inhibitors (DMIs) in the European diversity panel. (A) Relative growth was determined by comparing the fungal growth in fungicide-treated medium to that in control medium after 7 days. The growth area for each strain in the fungicide condition is normalized to the control condition. Red circles indicate strains with 100% resistance, while black circles represent strains with 0% resistance. (B) Schematic representation of the estimation of half maximal effective concentrations (EC50). Fungal strains were exposed to varying fungicide concentrations, and optical density (OD) measurements were taken at 0 days and 6 days post-inoculation (dpi). OD values are used to calculate EC50 values based on a non-linear logistic regression. Heatmaps display pairwise Pearson’s correlation coefficients, illustrating associations between (C) EC50 values and (D) relative growth values among the different DMIs.

### Complex genetic architecture of resistance to DMIs

We mapped mutations underpinning DMI resistance using GWAS. The DMIs prothiconazole, cyproconazole, prochloraz, tebuconazole, and epoxiconazole were assayed both on liquid (EC_50_) and solid medium. The recently approved mefentrifluconazole was only assayed for solid media growth. We investigated relative contributions of different polymorphism classes by performing separate GWAS for three different genotyping datasets; (1) SNPs, (2) large structural variation (variants >30 bp) and (3) 25-bp k-mer sequences derived from the short-read datasets uniquely mapped to a locus. We identified a total 21,220 genetic variants associated with DMI resistance, of which 12,557 (59%) overlapped with protein-coding regions across 158 genes (Figure 3B). The target gene *Cyp51* was universally mapped through associations with variation in resistance to all tested fungicides. Genetic variants near the multidrug resistance factor *MFS1* were associated with resistance to mefentrifluconazole, cyproconazole and epoxiconazole (Figure 3B). Discovery of genetic variation associated with DMI resistance was dependent on the phenotyping assay with 8,145 significant associations for EC_50_-based assays and 13,075 significant associations for relative growth measurements (Table S3). For epoxiconazole resistance, we detected 297 significant k-mers, 10 SNPs, and 1 structural variant based on the EC_50_ assays, whereas solid medium assays identified 5,412 k-mers, 156 SNPs, and 3 structural variants (Figure 3A). This discrepancy of associated variants may reflect differences in assay sensitivity and dose response properties to different DMI fungicides.

**Figure 3.**
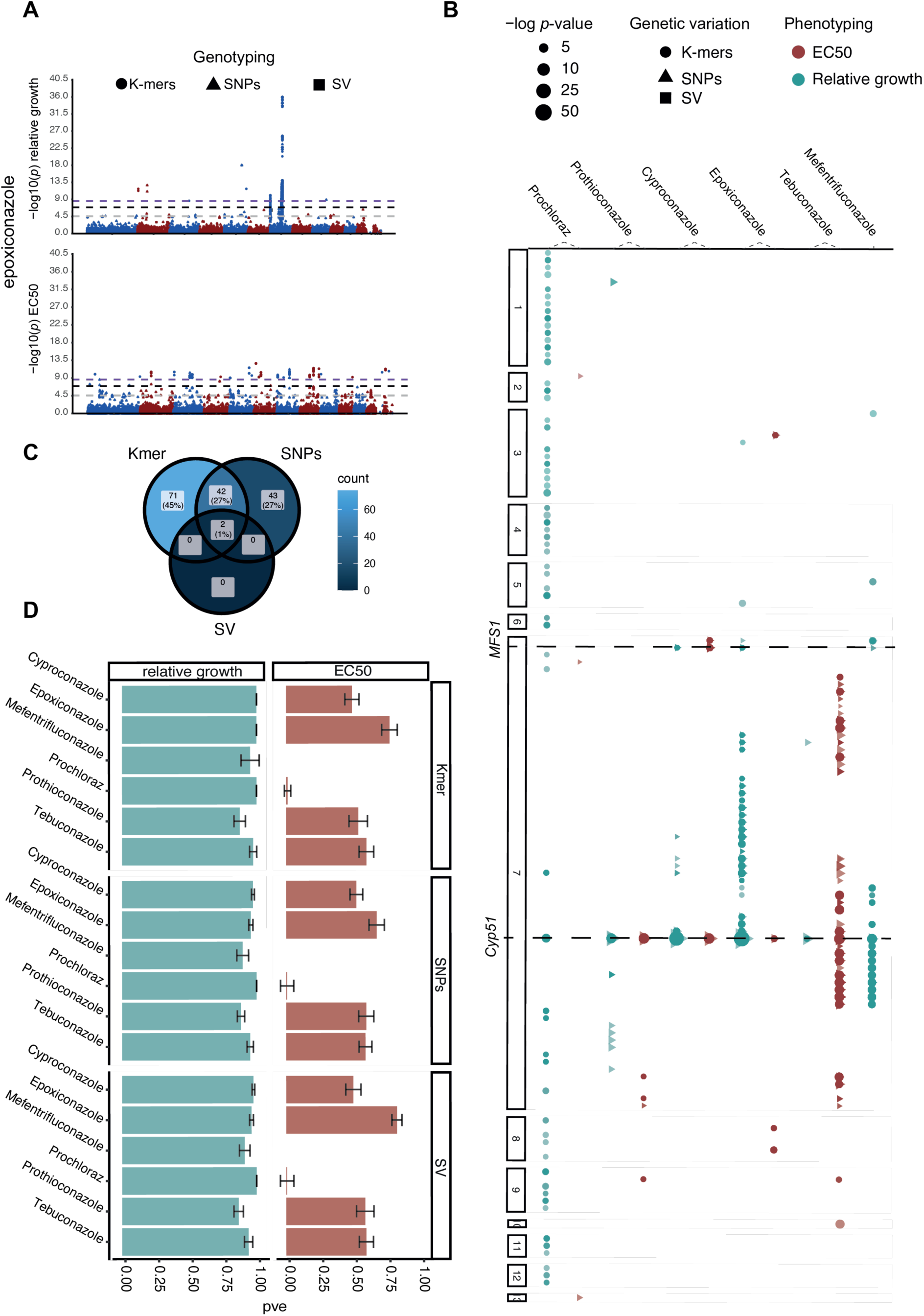
Genetic architecture of DMI resistance detected across the European diversity panel. (A) Manhattan plot of epoxiconazole resistance based on relative growth values (top) and EC50 estimates (bottom). For *p-*values (Wald) values >0.0001, we retained only 1 out of 30 positions for visualization purposes. (B) Scatter plot of the *p-*value (Wald) crossing the Bonferroni threshold overlapping with a gene across the 6 DMIs tested based on relative growth and EC50 values. Shape of the variants correspond to SNPs (dots), k-mers (triangle) and structural variation (square). (C) Venn diagram of genes associated with resistance representing overlaps between SNPs, k-mers and SVs. (D) Box plot of the chip heritability for SNPs, k-mers and SVs across the 6 DMIs tested based on relative growth and EC50 values.

Among the conditionally associated markers, we found 12 SVs, 20,274 k-mers and 934 SNPs significantly associated with prochloraz, mefentrifluconazole, epoxiconazole, prochloraz, cyproconazole and tebuconazole resistance. We found 12 SVs significantly associated with prochloraz, epoxiconazole, prochloraz, cyproconazole, tebuconazole and mefentrifluconazole resistance in relative growth and EC_50_-based phenotypes. An SV located 1387 bp downstream of *Cyp51* was associated with mefentrifluconazole, epoxiconazole, tebuconazole and cyproconazole resistance. Epoxiconazole resistance revealed two significant SV associations and 5412 k-mer associations in relative growth medium, associations with tebuconazole in EC_50_-based assays and mefentrifluconazole in relative growth medium, revealed substantial diversity, with 6,720 and 6,485 significant k-mers identified, respectively, compared to only 109 and 560 associated SNPs (Figure S4A). Resistance to prochloraz showed no significantly associated SVs in any of the two assays (Bonferroni threshold, α=0.05) but 366 significantly associated k-mers in relative growth on solid medium and 18 k-mers and three SNPs based EC_50_ measurements (Table S3A). Variation in the types of associated loci for the same assay may stem from challenges in robustly genotyping all genetic variants. However, variation in associated loci between assay types for the same DMI is most likely explained by distinct genetic contributions to colony growth or growth in liquid medium under fungicide stress.

To formally assess convergence in resistance mechanisms beyond the target genes, we focused on the 44 genes with significant associations for multiple fungicides or significant associations shared between both phenotyping assays (Figure S4B). We found that 45% (71/158) of the genes were uniquely associated with k-mer variants, and 27% (43/158) were uniquely associated with SNPs (Figure 4C). A total of 27% (42/158) genes were identified both by SNP and k-mer based associations with only 2% (2/158) of the genes being associated with all three genotyping methods: gene Zt09_7_00435 encoding a NAD(P)-binding domain superfamily protein and the gene Zt09_7_00444 serving unknown functions.

**Figure 4.**
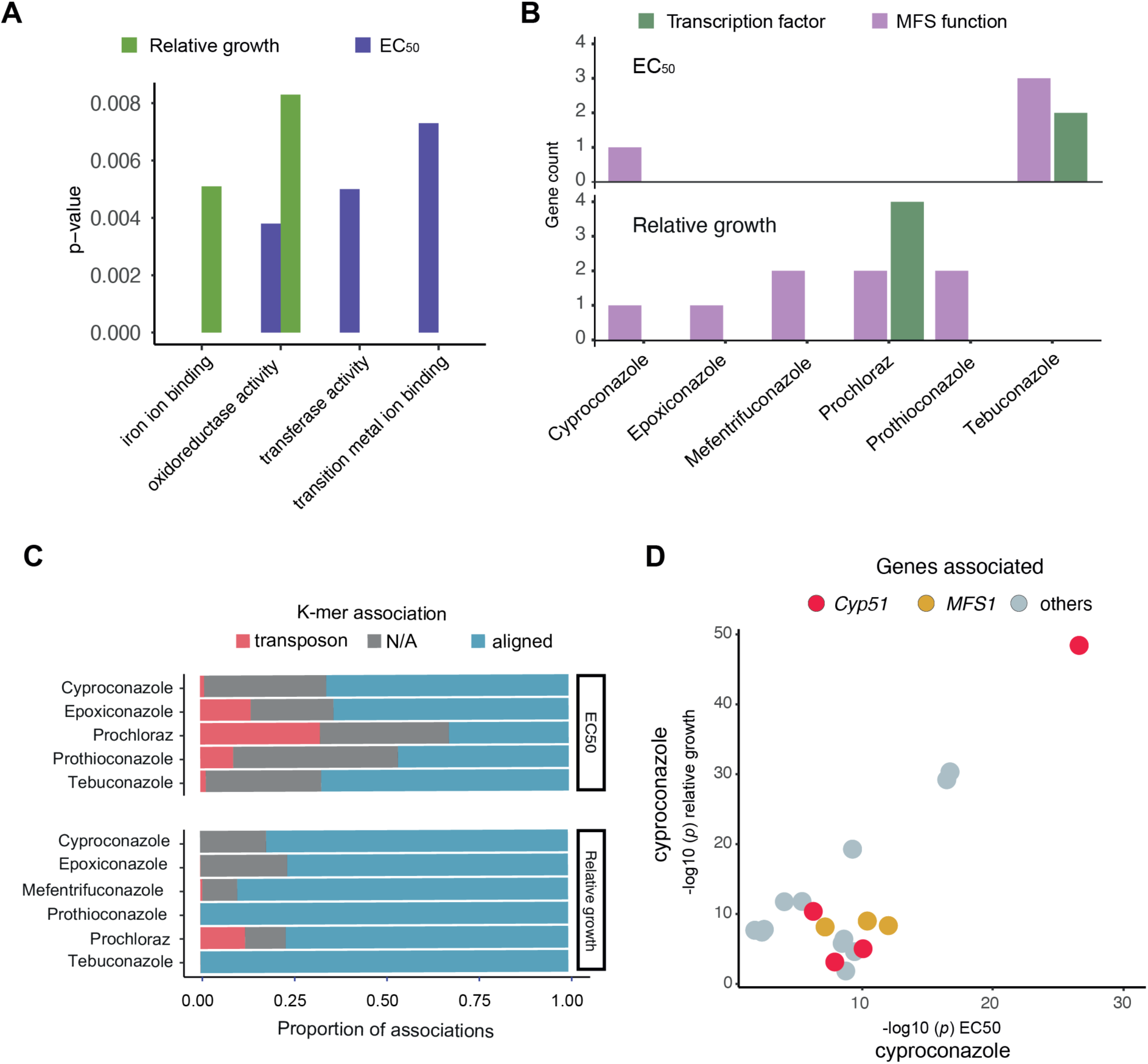
Transcription factors and MFS as resistance factors against DMIs. Enrichment for gene ontology (GO) terms. MFS and transcription factors were identified according to MFS (GO:0055085 - GO:0016021) and transcription factor (GO:0006355), respectively. (B) Enrichment analysis of the gene functions identified through GWAS based on EC50 and relative growth. (C) Percentage of k-mers aligning to the reference genome (red), to transposable elements (TEs) (blue) and not aligning (grey). Dot plot of the SNPs associated based on *p*-values based on relative growth and EC50 estimates.

To determine whether alternative genotyping strategies based on SNPs, k-mers or SVs facilitate GWAS discovery, we estimated the heritability of associated variants for each DMI and assay type. We found that k-mer-based associations explained overall a higher proportion of variation in resistance followed by SNPs and SVs, suggesting that k-mer variants capture more comprehensively genetic variation underlying resistance (Figure 4D). The heritability estimates for k-mer associations among DMIs and phenotyping assays ranged from 0.01-0.99, with an average of 0.96 for relative growth on solid medium and 0.48 for EC_50_ measurements. SNP-based associations explained heritability estimates of 0.944 for solid medium and 0.477 for EC_50_. SV-based associations explained heritability of 0.942 for solid medium and 0.52 for EC_50_ (Figure 3).

### Gene functions associated with DMI resistance

We identified a broad range of molecular functions contributing to DMI resistance. We performed an enrichment analysis on associated gene functions and found ubiquitin functions, acetyltransferase and amino acetyltransferase activity enriched for solid medium growth assays (Figure 4A). In contrast, gene functions discovered through liquid EC_50_ assays identified functions associated with oxidoreductase activity, transferase activity and transition metal ion binding, with the latter being consistently enriched independent of the assay type (Figure S5). Resistance to DMIs in *Z. tritici* and other filamentous fungi has been associated with efflux pumps (*i.e. MFS1*) (6, 60) and various transcription factors (8, 61). Our analysis identified a total of nine associations near *MFS* genes including *MFS1* based on solid growth assays (Figure S5). MFS transporters are often capable of exporting a diversity of compounds through cell membranes (62). Notably, our GWAS identified that associations in *MFS* loci are consistent with cross-resistance levels described in previous studies investigating MFS functions (57) (Figure S4A). Our GWAS also identified seven transcription factors associated with DMI resistance (Figure 4B). For prochloraz resistance alone, four transcription factors are contributing significantly to variation (genes Zt09_1_02033, Zt09_2_01091, Zt09_5_00447 and Zt09_7_00535).

Finally, we compared the strength of SNP-based associations with DMI resistance across the two phenotyping methods, *i.e.* EC_50_ values and relative growth. Our analysis revealed that the most significant associations varied depending on the phenotyping method with some variants showing strong associations with either EC_50_ or solid medium growth but not in both assay types. Notably, SNP associations in the *Cyp51* gene were significant in both assay types at the Bonferroni threshold, indicating that *Cyp51* variants contribute substantially to DMI resistance regardless of the growth conditions (Figure 4D). In conjunction, we demonstrate that DMI resistance across Europe has arisen from a combination of target site mutations in *Cyp51* as well as a wide range of non-target site mechanisms. Across all tested fungicides, *Cyp51* consistently appeared as the key genetic locus linked to resistance, highlighting the central role of this gene encoding the molecular target of DMI fungicides. Additional notable contributions are from transposable element polymorphism highlighting the complex genetic landscape of DMI resistance in Europe (Figure 4C).

### *Cyp51*-driven resistance gains

We further examined the contributions of *Cyp51* alleles on DMI resistance across Europe focusing on solid media growth (Figure 5A). We identified 18 significantly associated SNPs with 17 located in the coding sequence (CDS) and one in the intron (Figure S6A). Our GWAS confirms associations of previously documented missense resistance mutations (54, 63). For instance, the Ser524Thr variant showed association *p* = 10⁻⁵⁵ for epoxiconazole, *p* = 10⁻⁴⁸ for cyproconazole, and *p* =10^-29^ for prothioconazole. The Val136Phe mutation showed a *p* = 10⁻²² for epoxiconazole and *p* = 10^-10^ for cyproconazole. Additionally, 841 significant k-mers overlapped the *Cyp51* CDS.

**Figure 5.**
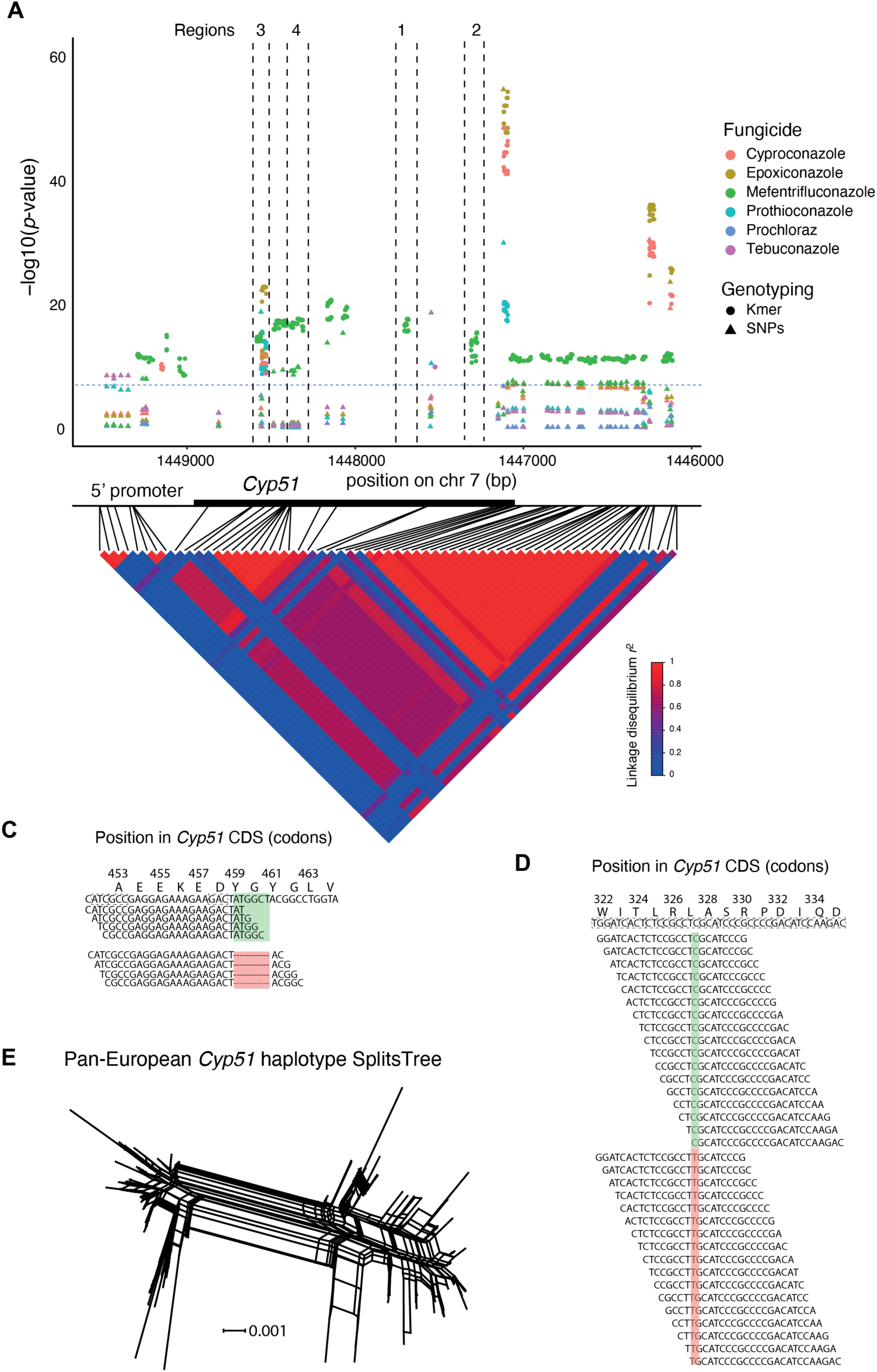
Complexity of *Cyp51* locus associations by SNP and kmer variants. (A) Manhattan plot of SNPs (dots), k-mers (triangle) associated with resistance to six DMIs. The black horizontal line shows the average SNP Bonferroni threshold (-log10 = 6.9) across the tested DMIs. The coordinates show positions in *Cyp51* and the linkage disequilibrium heatmap. (B) K-mers significantly associated to mefentrifluconazole aligning on the reference (IPO323) in the region stretching from 1447288 (bps) to 1447328 (bps) (Zone 1) which includes the two indels corresponding to Tyr459-Gly460Δ. Green shows k-mers positively associated with susceptibility, red shows positive associations to resistance. (C) K-mers significantly associated with mefentrifluconazole mapped from 1,447,659-7,700 correspond to the non-synonymous variant Leu327Leu (in zone 2). Green shows variants pos associated to susceptibility, in red significantly associated to resistance. (E) Pan-European phylogenetic network of the *Cyp51* gene haplotypes constructed using SplitsTree.

We examined associations of k-mers mapped to the *Cyp51* locus by focusing on four distinct regions showing independent associations compared to the SNP-based GWAS. Region 1 (k-mers 1,447,308-1,447,311) overlaping with a previously characterized indel polymorphism at codons Tyr459 and Gly460 associated with mefentrifluconazole resistance. We identified eight significantly associated k-mers with four aligning without mismatch to the reference IPO323 sequence of *Cyp51*, which represents a susceptible strain. As expected, the four k-mers were significantly associated with higher susceptibility. The remaining four k-mers aligned to the same region with one or more mismatches (Figure 5 C). In region 2 of *Cyp51* (k-mers 1,447,683-1,447,699), we identified 34 k-mers, with 17 of these associated with resistance to mefentrifluconazole and matching variation at a single synonymous SNP in the codon Leu327 (Figure 5D). 3 k-mers in region 3 (k-mers 1,448,572-1,448,574) were aligning to the reference sequence of the susceptible strain IPO323 and 3 k-mers aligned to the identical positions with a single point mutation corresponding to the synonymous SNP at Pro125 (Figure S7A). Finally, we found 73 k-mers matching region 4 (k-mers 1,448,318-1,448,359) each representing one or multiple synonymous variants at codons Ser158, Phe174, Arg177, Asn178, Asn179, His181, Thr187, Ser188 and Gly189 (Figure S7B). None of these synonymous variants were previously reported to be associated to fungicide resistance. SNP-based genotyping was unable to identify most of these associations due to the stringent quality-filtering applied to SNP variants (see Table S4).

To better understand the convergence in predicted effects of *Cyp51* mutations on DMI resistance, we investigated complements of missense and synonymous mutations for each DMI. We found that each DMI presented a unique combination of significantly associated variants. The smallest overlap in missense resistance variants was for tebuconazole against other DMIs with only the Ile381Val mutation being shared. In contrast, the missense mutation Ser524Thr was associated with prothioconazole, cyproconazole and epoxiconazole resistance. We also determined the resistance for 44 haplotypes representing the combinations of the most frequent 16 synonymous and missense mutations associated with DMI resistance (Table S5). The large number of synonymous and missense mutations in *Cyp51* underpins a highly diverse set of haplotypes across the European continent. This is illustrated by the complex phylogenetic network of *Cyp51* haplotypes (Figure 5E). As expected for the highly recombining populations of *Z. tritici, Cyp51* haplotypes show substantial diversification and significant evidence for recombination (*p* < 0.0001).

### Effects of *Cyp51* haplotypes on DMI resistance

Missense mutations in *Cyp51* are known to significantly affect DMI resistance. Here, we tested for effects of haplotypes carrying sets of significantly associated synonymous mutations. Using allele-swapping, we formally tested for the contribution of either sets of missense or synonymous mutations on DMI resistance (Table S6; Figure S6A). We selected the *Cyp*51 haplotype of strain 15STIRL021.1 collected in Ireland in 2015. Strains carrying this haplotype display strong resistance towards DMIs and carry a large number of resistance mutations (Table S1 and S5; haplotype number 40). The *Cyp51* haplotype of 15STIRL021.1 includes four missense mutations (Val136Cys, Ser188Asn, Ile381Val, Ser524Thr) and an indel resulting in the deletion of two amino acids (Tyr459-Gly460Δ). Besides missense mutations, the 15STIRL021.1 haplotype includes 14 synonymous mutations, which were associated with mefentrifluconazole resistance based on k-mer and SNP analyses (Fig 6A, Figure S7 A/B). To determine the impact of *Cyp*51 haplotypes experimentally, we swapped the wild type copy of the IPO323 *Cyp*51 haplotype with either one of the three following haplotypes, each under the control of the same tetracycline-repressible promoter: (1) the 15STIRL021.1 *Cyp51* haplotype including all synonymous and missense mutations (OE_IRL21), (2) the 15STIRL021.1 *Cyp*51 haplotype carrying all non-synonymous mutations but conserving original IPO323 codons for the 14 resistance-associated synonymous SNPs (OE_IRL21SYN), and (3) a reintegration of the IPO323 haplotype as a control (OE_IPO323) (Figure 6A).

**Figure 6.**
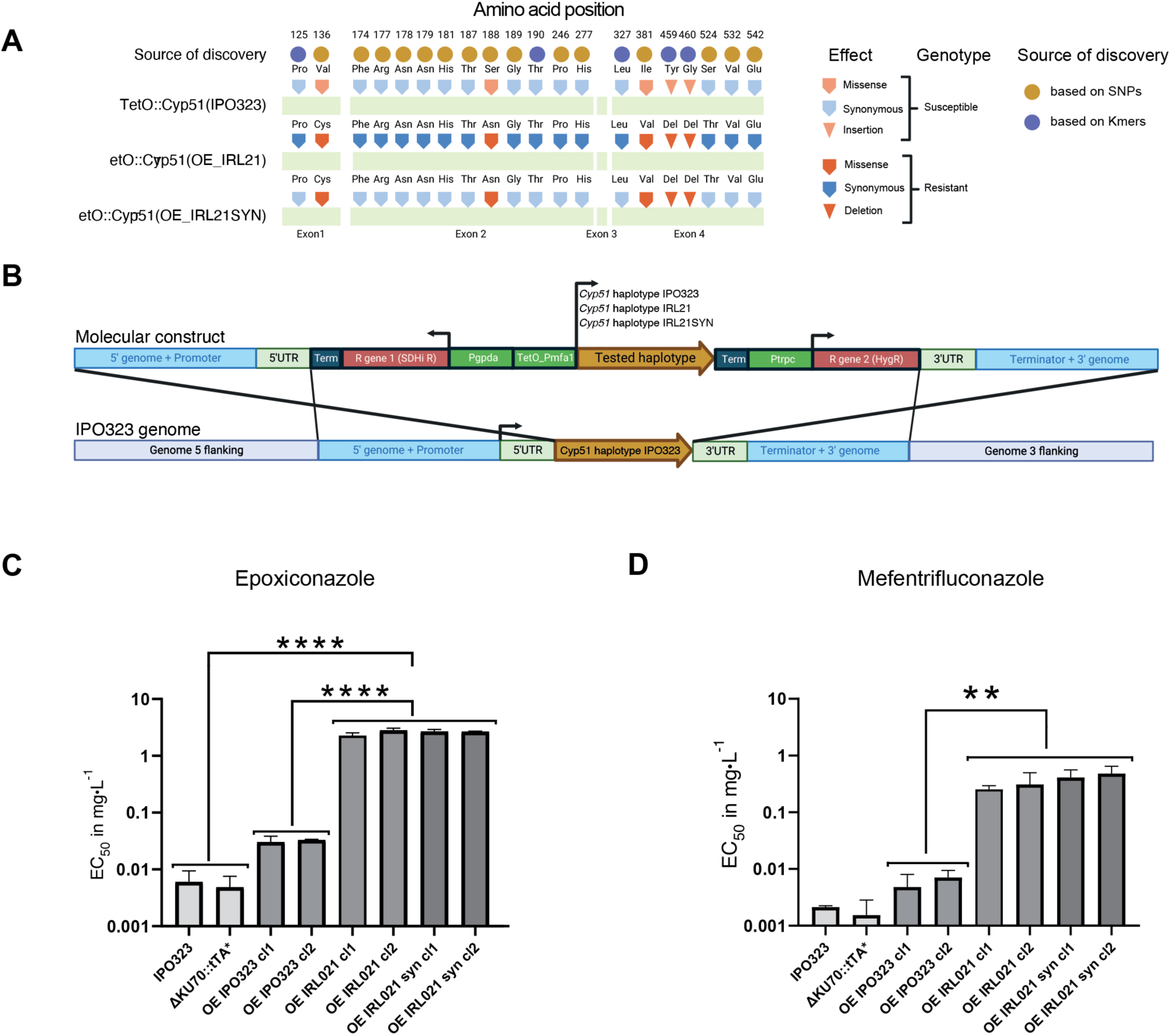
Assessment of *Cyp51* haplotype effects on DMI resistance. (A) Schematic of the susceptible and resistant *Cyp51* haplotypes of IPO323 (wild-type), the Irish strain 15STIRL21.1 (IRL21) and synonymous variants-only of the 15STIRL21 strain (SYN_IRL21). The top row shows mutated codon positions across *Cyp51* exons (light green bars). Filled light brown and dark blue circles report the GWAS detection method used to identify significantly associated variants. The filled arrows and triangles describe variant types: susceptible synonymous in light blue, synonymous missense in light orange, and matching resistance-associated synonymous variants in blue and resistance-associated missense and indel variants in orange . Triangles distinguish indels from SNPs (arrows). (B) Construct diagram of the *Cyp51* haplotype swapping in the IPO323 ΔKU70::tTA* mutant background. The center of the construct carries the to be tested haplotype of *Cyp51* (light brown) flanked by a tetracycline-repressible promoter (TetO_Pmfa1) in 5’ and by a terminator in 3’. This *Cyp51* expression cassette is itself flanked by two fungicide resistance cassettes (shown in red): isofetamid resistance and hygromycine resistance to exclude incomplete homologous recombination events during strain selection. Finally, 5’ and 3’ *Cyp51* flanking regions of the IPO323 genome were included for *in locus* homologous recombination. (C and D) Bar charts representing liquid culture EC50 values of the strains IPO323, ΔKU70::tTA*, OE_IPO323 (overexpression of IPO323 haplotype), OE_IRL21 (overexpression of IPO323 haplotype) and OE_IRL21SYN with epoxiconazole and mefentrifluconazole. One-way ANOVA: **** *p*-value < 0.0001, ** *p*-value < 0.05.

We assessed DMI resistance of the *Cyp*51-swapped IPO323 mutants with epoxiconazole and mefentrifluconazole (Figure 6B-C). On epoxiconazole, we detected a 80-fold difference in sensitivity between the control carrying *Cyp*51 IPO323 haplotype and its counterpart carrying the 15STIRL021.1 haplotype (average EC_50_ of 0.031 versus 2.539 mg.L-1, respectively), confirming a strong effect of this resistant haplotype. However, no significant difference was observed between the 15STIRL021.1 haplotype and its synthetic counterpart devoid of synonymous mutations (average EC_50_ of 2.539 mg.L-1 and 2.673 mg.L-1, respectively). We observed a similar resistance profile with mefentrifluconazole, with a 47-fold sensitivity difference between the control carrying *Cyp*51 IPO323 haplotype and its counterpart carrying the 15STIRL021.1 haplotype (average EC_50_ of 0.005 mg.L-1 versus 0.281 mg.L-1, respectively) and no consistent differences between 15STIRL021.1 and its synthetic analog (average EC_50_ of 0.281 mg.L-1 and 0.446 mg.L-1, respectively; Table S7). These results suggest that synonymous SNPs of *Cyp51* do not significantly contribute to resistance. The IPO323 control mutant showed 3.88 and 6.52-fold EC_50_ difference compared to the background ΔKU70::tTA* (EC_50_ of 0.031 and 0.005, respectively) in epoxiconazole and mefentrifluconazole (ANOVA; *p* > 0.05). This could be explained by the weak overexpression of the target under the control of the Tet promoter. Since a slight transcriptional overexpression of *Cyp*51 might mask a weak effect of synonymous mutations on protein translation efficiency, we tested the sensitivity of the mutants downregulated for *Cyp51* expression. As expected, transcriptional repression of the molecular target with doxycycline sensitized all mutants carrying the gene under transcriptional control of the Tet promoter. The mutants carrying the IPO323 wild type haplotype were strongly inhibited at the lowest epoxiconazole concentration in the test (0.00038 mg.L-1) and the mutant carrying the 15STIRL021.1 haplotype and its synthetic counterpart, showed approximately 15-fold higher sensitivity to epoxiconazole in the presence of doxycycline (EC_50_ of 0.173 and 2.539 mg.L-1 for 15STIRL021.1 haplotype, respectively). However, the addition of doxycycline did not modify the lack of sensitivity differences between the 15STIRL021.1 haplotype and its synthetic counterpart lacking synonymous mutations (EC50 of 0.173 and 0.266 mg.L-1 respectively, Figure S8A). As a control, we used benzovindiflupyr (an SDHI) and observed no significant differences among the IPO323 mutant strains (see Figure S8B-C). Hence, the assay did establish strong effects of the 15STIRL021.1 *Cyp51* haplotype compared to a sensitive strain. Even though synonymous mutations showed strong associations in GWAS, our assay confirmed only contributions by missense mutations to DMI resistance.

## Discussion

We established a large strain panel for the major wheat pathogen *Z. tritici* to capture recent gains in fungicide resistance. The European diversity panel covers distinct agricultural regions with variable intensities of recent DMI fungicide application. Our genome sequencing analyses revealed that Europe harbors a highly diverse set of genotypes central to our understanding of *Z. tritici* diversity globally. We assessed two complementary sensitivity screening method, revealing notable variation in resistance profiles across time and geography. We detected a total of 21,220 variants in 158 genes associated with resistance to 11 DMIs. The scope of the associations across the genome allowed for fine-grained analyses of cross-resistance among the structurally related DMIs, as well as unravel patterns of parallel resistance evolution across the European continent.

The European diversity panel revealed extensive admixture across the continent with only weak genetic differentiation (*i.e.* isolation-by-distance) over large distances. Exceptions included well differentiated genetic groups sampled in Southern Europe (*i.e.*, Italy, Spain, and Southern France) and Eastern Europe (*<u>i.e.</u>*, Russia), which likely reflects recent gene flow from North Africa and Central Asia, respectively (26). The substantial genetic diversity and weak differentiation is expected to empower GWAS by reducing spurious marker associations (64). Similarly, resolution of GWAS is improved by reproducible phenotyping methods capturing traits relevant for resistance gains in the field. Leaf pathogens such as *Z. tritici* are usually exposed to fungicides during early disease development on the host. Fungicides act on spore germination stage or at later mycelial development stages prior to full infections (65). DMIs are known to inhibit fungal mycelial development stages (66). Our resistance assays were performed using either liquid culture or colony growth measurements. Even though reproducibility of each assay was high, resistance captured by each of the methods was only weakly correlated. This suggests that the pathogen expresses DMI resistance through at least partially distinct mechanisms in these two conditions. Furthermore, differences in bioavailability of fungicides in liquid cultures compared to solid agar may explain additional discrepancies. For example, reduced oxygen levels in liquid culture and other microenvironmental factors may impact variation in the expression of resistance. The largest discrepancies between assays were found for prochloraz and tebuconazole. Loci mapping power may have been weakly affected by differences in sample size for the different fungicide assays. In addition, cryptic switches in growth morphology (yeast or filamentous growth) may also account for some of the variation (67). The dual assays in liquid culture and on solid medium may potentially better capture the broader range of resistance mechanisms evolved across the numerous conditions experienced during the pathogen life cycle.

### Genetic architecture of DMI resistance across Europe

We identified a total 21,220 genetic variants associated with DMI resistance, of which 12,557 (59 %) were in protein-coding regions covering 158 distinct genes. Recent work revealed that a significant portion of DMI resistance is encoded by structural variation beyond SNPs (6, 17, 57). To map such genetic variations beyond point mutations, we implemented genotyping approaches suitable for the robust discovery of complex variants. We controlled for spurious genotypes by integrating reference-genome based controls for variants assessed from short read data. Structural variants were identified based on a pangenome graph approach and subread associations (*i.e.* k-mers) were cross-checked with reference genome mapping. We found a very small (2%) overlap among gene sets mapped by the three different genotyping approaches. K-mer and SNP-based approaches revealed a shared gene set of 27% of the total. Hence, our approach of combining multiple independent genotyping approaches has significantly expanded the scope of mapped resistance variation.

The gene encoding the DMI target Cyp51 was well-resolved by GWAS with 18 SNPs and 841 k-mers mapped to the coding sequence further confirming that mutations in the target are the primary driver of DMI resistance in the field (50, 51, 54, 55). Similarly, the gene encoding the detoxification factor MFS1 was mapped by four variants (1 SNP and 3 k-mers). Previous work established that multidrug resistance by MFS1 is modulated by regulatory elements upstream of the gene (6, 57). We confirmed that MFS1 is likely contributing to DMI resistance through regulatory tuning by identifying 75 mapped k-mers upstream of the gene. These k-mers are likely capturing TE insertion variation modulating gene expression such as the previously identified TE insertion 519 bp upstream associated to prochloraz, epoxiconazole and prochloraz sensitivity shifts (6). Neither *Cyp51* or *MFS1* were amenable to SNPs based GWAS discovery in previous studies on *Z. tritici* and their implication in resistance was identified previously only through mutant analyses and crosses (6, 54). Mapping *Cyp51* and *MFS1* is consistent with the substantial increase in mapping power of the European diversity panel. We note though that we were unable to replicate an association of a DHHC palmitoyltransferase locus with propiconazole resistance (17). The geographic scope of the palmitoyltransferase association was restricted to a North American population though, suggesting a lack in convergence on this mechanism in Europe. Finally, even though the ABC transporter gene *Atr1* and its promoter were shown to contribute to cyproconazole resistance in mutants (68), no genetic variants were found to be significantly associated. We cannot rule out the involvement of Atr1 overexpression in our panel since one of the mapped transcription factor might indeed drive Atr1 expression, similar to how the transcription factor Mrr1 regulates the ABC transporter AtrB in *Botrytis cinerea* (69, 70). Lack of association in ABC transporters or promoters thereof may also be due to differences in the genetic background of the strains, variation in experimental conditions to assess DMI resistance or that the European diversity panel did not capture the relevant mutations at high enough frequency (detection limit of 5% in SNPs).

In addition to the effective mapping of variations driving DMI resistance at known loci, we indentified a broader set of previously undescribed resistance factors. These included seven genes encoding potential drug export functions by transporters belonging to the major facilitator superfamily (MFS) beyond MFS1, and seven transcription factors. The associations in MFS genes were narrower across the spectrum of DMIs. This may be due to the variation in chemical structures among DMIs and possible links to specific fungicide-exporting capabilities by each MFS. If MFS transporters confer resistance to specific DMI subclasses, fungicide deployment might be optimized to account for previously unknown standing MFS-associated resistance in the field. Transcription factors modulating azole resistance have been supported by molecular characterizations in *e.g. B. cinerea* and *Saccharomyces cerevisiae* (61, 71, 72). We mapped four transcription factors associated with prochloraz resistance in solid media relative growth conditions and three transcription factors associated with tebuconazole resistance based on EC_50_. In previous work, transcription factors were often indirectly linked to DMI resistance, because they tend to regulate multiple genes, challenging the establishment of causal links with DMI resistance. The discovery of transcription factors associated with DMI resistance underpins the central role of transcriptional reprogramming in mediating fungicide resistance in very different fungal pathogens (70, 73) .

### Emergence of resistant *Cyp51* haplotypes

The role of missense mutations in *Cyp51* has been extensively assessed experimentally, whether synonymous mutations can impact resistance remains unknown (74). Here, we discovered numerous significant associations of both synonymous and missense mutations in *Cyp51*. GWAS linked missense variants to previously functionally validated alleles. We found that Ile381Val and Ser524Thr were associated to prothioconazole and epoxiconazole resistance, respectively. We also found discrepancies in variants associated with prochloraz resistance. For instance, we found no evidence for an involvement of Ser524Thr even though previous mutant analyses demonstrated an effect (54). These discrepancies may be due to epistatic interactions among *Cyp51* mutations, where the combined effects of multiple mutations create unique resistance profiles that vary depending on the specific DMI tested. Epistatic effects of *Z. tritici Cyp51* mutations were demonstrated using functional replacement in yeast (54). The combination of Leu50Ser and Tyr461Ser missense mutations conferred an increase in resistance by a factor 6 to epoxiconazole and a factor of 52 to tebuconazole. Adding the Ser524Thr mutation to the two previous mutations increases the resistance factor to 95 for epoxiconazole and 632 for tebuconazole. Furthermore, tested in isolation some mutations produced no functional Cyp51 highlighting co-dependencies of resistance-conferring mutations in both conferring resistance and maintaining enzymatic function. Beyond epistatic effects of resistance-conferring mutations, GWAS is sensitive to LD in mapping populations. Strong recent selection of resistant *Cyp51* haplotypes is expected to emphasize the association of individual mutations at short physical distance. Combining GWAS discovery with specific genetic validation of associated haplotypes will alleviate these constraints on mapping power. In this study, we tested whether synonymous *Cyp51* mutations which accumulated during resistance gains on the European continent could contribute to resistance. For this, we introduced either the full complement of gained synonymous and missense *Cyp51* mutations into a sensitive background or only missense mutations. The full exchange for a resistant *Cyp51* haplotype significantly increased resistance consistent with previous findings, however no significant contribution of synonymous mutations was detectable, suggesting these associations were due to LD. Looking ahead, our gene swapping approach can be more broadly applied to reveal epistasis and decipher further complex genetic contributions in the most polymorphic loci.

In conclusion, our work establishes a highly diverse panel of field-collected strains of a crop pathogen capturing gains in DMI resistance at the level of the European continent. The large body of associated variants across diverse gene functions highlights the complexity of resistance gains in agriculture. Managing resistance remains a challenge for agricultural production but genomic monitoring opens powerful perspectives on this process. Development of new compounds may benefit from understanding the genetic mechanisms underlying resistance and facilitating the targeting of novel pathways. Additionally, integrating field-specific resistance surveillance with predictive genomic tools will enable more effective and sustainable fungicide deployment strategies.

## Methods

### European diversity panel

For resistance monitoring purposes, Syngenta sampled pathogens from commercial and trial sites across Europe. The fungicide resistance monitoring efforts for *Z. tritici* led to a collection of 8607 strains. From this collection, we defined a representative subset to conduct this study on DMI resistance. Strains were selected from the collection based on a hierarchical clustering approach grouping sampling locations into 100-km radius areas. Within each area, we balanced the collection over the 2005-2019 sampling years based on availability. The assayed European diversity panel included 1394 strains from 27 different countries. A total of 286 strains were previously sequenced (26) (Table S1) and 1134 were sequenced for this study. The European diversity panel was contrasted against an additional 736 strains from a global diversity survey of the pathogen (26) (Table S1).

### Purification of the European diversity panel

All strains included in the European diversity panel were collected from infected leaves across Europe, preferably with visible pycnidia, were wrapped in dry paper towels and packed into a paper envelope for shipment. Upon arrival in the laboratory, samples were labeled with a unique code and the arrival date was recorded. The leaves were dried, wrapped in fresh paper towels if necessary, and stored at 4°C until further processing. Leaves with symptoms were cut into 2 cm pieces, surface-sterilized in 2% bleach for 2 minutes, and rinsed with sterile distilled water. Leaf cuts were placed on wet filter paper in Petri dishes (1.3 mL water for 9 cm dishes) and incubated at 20°C for 24 hours. Single strains were picked from cirrhi under a binocular microscope and transferred to V8 agar plates with antibiotics. Plates were incubated for 4-7 days at 20°C. To ensure that only a single genotype was collected per sample, a single spore isolation step was performed for every sample. Single colonies were subcultured on fresh V8 plates and incubated under the same conditions for an additional 7 days. Each strain was stored independently in cryovial preserved in liquid nitrogen. Fresh cells harvested from V8 plates incubated at 18°C for a period of five days were used as inoculum for all experiments. Media recipes used in this study are summarized in Table S8.

### Fungicide sensitivity assays

Microtiter plate liquid fungicide resistance assays were conducted to determine the half maximal effective concentration (EC_50_) as follow. Spore suspensions were obtained from YPD plates incubated for 6 days, 20°C, in the dark and standardized to reach a density of 100’000 spores per ml using a Neubauer counting chamber. Fungicides, pre-dissolved in DMSO were added at 1% volume to liquid YBG-Medium carrying the spore suspension (100 µl) (75). For prothiconazole and cyproconazole, tested concentrations were obtained through a 4-fold cascade dilution series, starting from 40 mg·L⁻¹ and sequentially decreasing to 0. 0098 mg.L-1 (7 inhibitor concentrations + DMSO control), for prochloraz, epoxiconazole and tebuconazole starting from 10 and sequentially decreasing to 0.0024 mg.L-1 (7 inhibitor concentrations + DMSO control). Sterile 96 wells microtiter plates with lids (Costar) were incubated for 6 days at 20°C in the dark, measurements were performed at 405nm absorbance using a plate reader (Envision 2105, Perkin Elmer, Waltham, USA). EC_50_ values were calculated using AGSTAT developed by Syngenta using a nonlinear curve fit. EC_50_ values and details on all tested fungicides can be found in Table S1.

Fungicide sensitivity on solid medium was determined using relative colony size estimates (fixed fungicide concentration). The European diversity panel strains stored at -80C° were arrayed in 96-well cryo-stock plates in sterile solution carrying 8,5% skim milk (m/vol) with 10% glycerol (used as mother array plates). From the mother array plates, the panel of strains to be tested were transferred using a 96-floating pin replicator tool (408FS2AS, V&P Scientific Inc.) to 96-well flat bottom plates (model 3370, Corning) pre-filled with 100μl YPD liquid medium and left to grow at 18°C for 7 days to reach a growth plateau. Spore suspensions were not further standardized and directly spotted using a Rotor HDA (Rotor+) (Singer Inc., Watchet, UK) with sterile pins (RePads) (Singer Inc., Watchet, UK) onto AE agar media plates (PlusPlates) amended or not with fungicides. The fungicides were dissolved in DMSO and then mixed with AE-agar to achieve a final concentration of 1% DMSO. Five different fungicide concentrations were applied to achieve final concentrations of 50, 10, 1, 0.1, and 0.01 mg.L-1 per plate. Single concentrations were then selected for further GWAS studies. Control plates only contained AE-agar with 1% DMSO. Spotted AE-agar plates were incubated at 20°C for 7 days in the dark before imaging. Image capture was performed using a Phenobooth (Singer, Watchet, UK) and image analysis performed with the Phenosuite (version 2.21) software package (Singer Inc., Watchet,) by comparing paired growth areas of colonies grown on control plates without fungicide (DMSO controls) with fungicide-amended plates. The relative growth area on amended media (*area*_e_) relative to controls (*area*_c_) was estimated using the following formula:

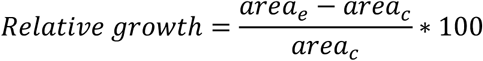

Plates with signs of contaminations were excluded. Plates with >2x more growth on fungicide-amended plates *vs.* controls were excluded as well. Relative growth estimates and assayed fungicides are reported in Table S1 and Table S9.

Fungicide sensitivity of the transformants was determined in liquid culture assays. Sensitivity assays were performed as previously described with the following differences: Spore suspensions were obtained from YPD plates rather than YBG with identical incubation steps. For epoxiconazole, mefentrifluconazole and benzovindiflupyr (SDHI control), 11 concentrations and a DMSO control were tested for each fungicide. The assayed concentrations were obtained through a 4-fold cascade dilution series, starting from 40 mg·L⁻¹ and sequentially decreasing to 0.000038 mg.L-1. Growth measurements were performed at 600 nm absorbance using a plate reader (Envision 2105, Perkin Elmer, Waltham, USA). EC_50_ values were calculated using four-parameter non-linear curve fitting using the software GraphPad Prism 10. To assess the effect of *Cyp51* repression, 30 mg.L-1 of doxycycline (DOX) was added to the sensitivity assays.

### DNA extraction, Illumina sequencing, and SNP calling

Whole-genome sequencing data was produced from high-quality genomic DNA extracted using the DNeasy Plant Mini Kits (Qiagen Inc.) following the manufacturer’s instructions. Paired-end sequencing of 250 cycles with an ∼500 bp insert size was performed by Novogene Inc. using the Illumina NovaSeq 6000 platform. We used Trimmomatic v.0.39 to trim low-quality sequencing reads and remove adapter contamination in each strain (76). Filtered sequences were aligned to the *Z. tritici* reference genome IPO323 (43) using Bowtie2 v.2.3.3 (77). The Genome Analysis Toolkit (GATK) v.4.0.1.2 (78) was used for single nucleotide polymorphism (SNP) calling and variant filtration. The GATK HaplotypeCaller was run with the command -emitRefConfidence GVCF and -sample_ploidy 1. Joint variant calling was performed using the tool GenotypeGVCFs merging HaplotypeCaller gvcfs produced for an additional 1022 from a previous study (26) with the option -maxAlt 2.

### Population genetic analysis and linkage disequilibrium

The joint genotyping calls for the European diversity panel (*n* = 1420 strains before filtering) supplemented by strains from additional continents (total *n* = 2156 strains) were filtered using vcftools v. 0.1.16 (79). For population structure analyses, we thinned the SNP set to retain variants satisfying the following criteria: a single variant per 1000 bp (--thin 1000), a minor allele frequency (MAF) ≥0.05 (--maf 0.05) and less than 10% missing genotypes (--max-missing 0.9). This filtering retained 22,614 SNPs. We performed a principal component analysis (PCA) using the read.vcfR and the dudi.pca functions in R (80). To analyze genetic structure within the European diversity panel, the same filtering was applied to the 1420 strains retaining 22,710 SNPs. To perform maximum likelihood estimation of individual ancestries, we used Admixture v. 1.3.0 (81). We chose an admixture model independent of prior population information and with correlated allele frequencies. The admixture model was run with 10,000 iterations. We assessed K between 2-20, with 5 repetitions per K. The most likely number of populations (K) was estimated with the cross-validation method and output analyzed with the R package Pophelper (82). The genetic distance (*D_ST_*) between individuals was estimated using PLINK v1.90 (83). Finally, geographic distance between strains was calculated using the distHaversine function of the geosphere R package (84) (Table S10).

We analyzed the decay in LD on chromosome 1. For this, we used all SNPs with a minor allele frequency ≥5 %. We calculated the linkage disequilibrium *r^2^* between marker pairs using the option -- hap-r2 and --ld-window-bp 10000 in VCFtools v.0.1.15 (79) with. The decay of LD with physical distance was estimated using a non-linear regression model (85, 86).

### SNP-based genome-wide association mapping

For SNP-based GWAS on the Europe diversity panel, we retained SNPs with a genotyping call rate of ≥90% and a MAF ≥5% resulting in a final set of 472,103 biallelic SNPs (Table S1). Traits for GWAS included the EC_50_ estimates for tebuconazole, cyproconazole, prothioconazole, epoxiconazole, and prochloraz. For relative growth, we included the same compounds along with mefentrifluconazole (Table S2). We accounted for relatedness by constructing a genetic relatedness matrix (GRM) among strains using all genome-wide SNPs with the option “-gk 2” in GEMMA (87). Thus, all the associations were performed using a univariate linear mixed model (MLM+K) where K is the GRM as a random effect:

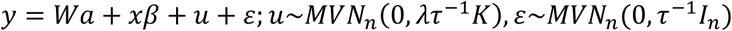

*y* represents a vector of phenotypic values for *n* individuals; *W* is a matrix of covariates (fixed effects with a column vector of 1), *α* is a vector of the corresponding coefficients, including the intercept; *x* is a vector of the genotypes of the SNP marker, *β* is the effect size of the marker; *u* is a vector of random individual effects; *ε* is a vector of random error; *τ*-1 is the variance of the residual errors; *λ* is the ratio between the two variance components; *K* is the *n* × *n* GRM and *I_n_* is an *n* × *n* identity matrix and *MVN_n_* represents the multivariate normal distribution. We applied a stringent Bonferroni threshold (*α* = 0.05; *p* = *α* / total number of SNPs) to identify most robust SNP associations. Significant SNPs were annotated using snpEff v5.0e (88). We provide all association mapping outcomes in Table S3.

### K-mer based genome-wide association mapping

We performed k-mer based GWAS on the same 11 traits used for SNP-based GWAS following a previously described approach (89). We used k-mers of 25-bp length as suggested for small genomes such as *Z. tritici* following previous studies (29). Quality-filtered sequencing reads (see above) of 1406 strains from the European diversity panel were used for k-mer screening. K-mers were counted using a two-step process: The canonization involved treating each k-mer and its reverse complement as equivalent. In contrast, non-canonization treats the k-mer and its reverse complement as distinct entities. A *k*-mer and the reverse-complement are supposed to have the same chance to appear in fastq files. However, technical reasons can create a bias to one of the forms for some *k*-mer. Therefore k-mers were then filtered based on two criteria: (1) a k-mer must appear in both its canonized and non-canonized forms in at least 5% of the strains, and (2) it must appear in both forms in at least 20% of the strains in which it was found.

A GRM was estimated with EMMA (Efficient Mixed-Model Association) as an identity-by-state (IBS) matrix under the assumption that each k-mer has a small, random effect on the phenotype. GWAS was performed by using an LMM+K model in GEMMA with a likelihood ratio test to determine *p*-values. Beta estimation and significance testing were performed using the patched version of kmers_gwas.py, incorporating the recommendations outlined (https://github.com/voichek/kmersGWAS/issues/53 and https://github.com/voichek/kmersGWAS/issues/91). A k-mer was significant when the *p*-value passed the permutation-based threshold as described by Voichek and Weigel (2020). We attempted to map all significant k-mers for each trait to the reference genome using the short-read aligner bowtie v1.2.2 (77) with the command “-a --best --strata” retaining the k-mers with a unique alignment. We used the center position of the mapped k-mer to the reference genome as a coordinate to inspect nearby features using BEDtools v2.31.0 (90). In total number of k-mers consisted of 1,159,029,650 unique k-mers. *Cyp51* sequences were used for the construction of an unrooted phylogenetic network with SplitsTree v4.17.1

(91). We provide links to all significant association mapping outcomes in Table S3.

### Structural variation-based genome-wide association mapping

We identified insertion and deletion variants ≥30bp in 18 reference-quality genomes (92) by aligning these to the IPO323 reference using MUM&Co (93) and minimap2+paftools (94). We used Jasmine (95) to combine the structural variant predictions from each genome into a single variant catalog with 25,839 non-overlapping sites based on IPO323 reference genome coordinates. The variant catalogue was used to build a pangenome graph with the *vg* toolkit (96). Graph variants were genotyped in all strains by mapping quality-trimmed Illumina reads to the graph using Giraffe (97). We set all genotype calls with fewer <4 supporting reads to missing and excluded all multi-allelic and invariant sites using GATK v4.2.41 (98). GWAS was performed on a subset of 1135 strains (Table S1) using the software GEMMA (87). We accounted for relatedness by including a GRM and filtered variants for MAF≥0.05. We provide data on all significant association mapping outcomes in Table S3.

### Gene functions and heritability

Genes located near significant variants were identified based on the functional annotation of the IPO323 genome (99) and using the “intersect” command in BEDtools v. 2.29.2 (90). Gene ontology (GO) terms for encoded proteins we determined using InterProScan v. 5.36–75.0 with default parameters (100). GO enrichment analyses were performed using the R packages GSEABase v. 1.35.0 and GOstats v. 2.38.1 (101). GO enrichment analyses were performed only for GO terms with a minimum term size of five genes and a false discovery rate (FDR) of 0.01 in hypergeometric tests. The proportion of variance in phenotypes explained (PVE) by genotypes, often referred to as ’chip heritability’, was estimated using the LMM implemented in the EMMA software (Zhou and Stephens, 2012) (Table S11). This approach allows for the inclusion of both fixed and random effects, thereby accounting for population structure and relatedness among individuals contributing to more robust estimates of heritability.

### Molecular methods for transformations

All plasmids used for transformation were derived from the previously described binary vector pNOV2114 (102). Inserts were designed *in silico* and synthesized by GENEWIZ Inc. (Germany). To functionally replace the genomic *Cyp51* IPO323 copy with doxycycline-repressible *Cyp51* haplotypes, we first generated a ΔKU70::TtA* strain carrying an expression cassette of the TtA* (103) at the KU70 locus (binary plasmid pNOV2114-Ku70_TetRep_NPT2_Ku70). We then designed three plasmids to replace *Cyp51 in-locus*: i) AltsdhC_Hygr_pNOV2114_CYP51_IPO323, representing the IPO323 haplotype, ii) AltsdhC_Hygr_pNOV2114_CYP51_IRL021 representing the haplotype of strain 15IRL021 and AltsdhC_Hygr_pNOV2114_CYP51_IRL021_SYN representing the haplotype of strain 15IRL021 but carrying all synonymous mutations matching the IPO323 haplotype (Table S12 & Table S13). The inserts obtained by gene synthesis were cloned between *EcoRI* and *HindIII* sites. To ensure full gene replacement, two different resistance cassettes were used placed upstream and downstream of *Cyp51*, respectively. The flanking region of 1135 bp upstream of *Cyp51* is followed by an isofetamid resistance cassette (PtrpC::alt-sdhC::MoILV term in reverse orientation), followed by a TetO promoter (Zarnack et al., 2006) driving tetracycline-repressible overexpression of *Cyp51* haplotypes. Downstream of the 3’ end of *Cyp51*, a hygromycin-resistant cassette was inserted, followed by 1042 bp corresponding to downstream genomic sequence (see figure 6A for graphical overview, and supplemental file Table S13 with plasmids sequences). *Agrobacterium tumefaciens* mediated transformations were carried out following established protocols (Bowler et al., 2010). for *Cyp51 in situ* replacement. The IPO323DKu70::TtA background was selected on G418 (250 mg.L-1 on AE agar media) and subsequent *Cyp51* swapping transformants were selected on combined hygromycin and isofetamid selection (in AE media, at concentrations of 100mg.L-1 and 10mg.L-1, respectively) . Clones were confirmed by PCR using primer combinations and PCR conditions listed in Table S14.

## Declarations

## Acknowledgements

We thank Nadja Lindenberger, Regula Frey, Salvatore Accardo and Syngenta’s monitoring team for collecting the strains analyzed in this work. We also thank Lucio Garcia and the NGS platform at Syngenta for assistance with sequencing. Sabina Tralamazza and Ursula Oggenfuss provided insightful discussions.

## Author contributions

GP, GS and DC conceived the study, GP performed research and analyzed data. GP, DF, DE and CD performed analyses. SFFT provided materials. EGT provided datasets. TB and AF curated datasets. GS and DC supervised the work and acquired funding. GP, DC and GS wrote the manuscript.

## Data availability

Genome sequencing data is available on the NCBI Sequence Read Archive (accession numbers are provided in Supplementary Table S1). Additional datasets are made available in supplementary materials and Zenodo (https://doi.org/10.5281/zenodo.15052606)

## References

1. Steinberg G, Gurr SJ. 2020. Fungi, fungicide discovery and global food security. Fungal Genet Biol 144:103476.

2. Corkley I, Fraaije B, Hawkins N. 2022. Fungicide resistance management: Maximizing the effective life of plant protection products. Plant Pathol 71:150–169.

3. Hawkins NJ. 2024. Assessing the predictability of fungicide resistance evolution through in vitro selection. J Plant Dis Prot 131:1257–1264.

4. Hawkins NJ, Bass C, Dixon A, Neve P. 2019. The evolutionary origins of pesticide resistance: The evolutionary origins of pesticide resistance. Biol Rev 94:135–155.

5. Fan F, Zhu Y-X, Wu M-Y, Yin W-X, Li G-Q, Hahn M, Hamada MS, Luo C-X. 2023. Mitochondrial Inner Membrane ABC Transporter Bcmdl1 Is Involved in Conidial Germination, Virulence, and Resistance to Anilinopyrimidine Fungicides in Botrytis cinerea. Microbiol Spectr e00108–23.

6. Omrane S, Sghyer H, Audéon C, Lanen C, Duplaix C, Walker A-S, Fillinger S. 2015. Fungicide efflux and the MgMFS1 transporter contribute to the multidrug resistance phenotype in *Z ymoseptoria tritici* field isolates: Fungicide efflux & *MgMFS 1* contribute to MDR in *Z. tritici*. Environ Microbiol 17:2805–2823.

7. Zhang T, Xu Q, Sun X, Li H. 2013. The calcineurin-responsive transcription factor Crz1 is required for conidation, full virulence and DMI resistance in Penicillium digitatum. Microbiol Res 168:211–222.

8. Shrivastava M, Kouyoumdjian GS, Kirbizakis E, Ruiz D, Henry M, Vincent AT, Sellam A, Whiteway M. 2023. The Adr1 transcription factor directs regulation of the ergosterol pathway and azole resistance in Candida albicans. mBio 14:e0180723.

9. Vicentini SNC, Moreira SI, Da Silva AG, De Oliveira TYK, Silva TC, Assis Junior FG, Krug LD, De Paiva Custódio AA, Leite Júnior RP, Teodoro PE, Fraaije B, Ceresini PC. 2022. Efflux Pumps and Multidrug-Resistance in Pyricularia oryzae Triticum Lineage. Agronomy 12:2068.

10. Celia-Sanchez BN, Mangum B, Brewer M, Momany M. 2022. Analysis of Cyp51 protein sequences shows 4 major Cyp51 gene family groups across Fungi. G3 GenesGenomesGenetics jkac249.

11. Zhang J, Li L, Lv Q, Yan L, Wang Y, Jiang Y. 2019. The Fungal CYP51s: Their Functions, Structures, Related Drug Resistance, and Inhibitors. Front Microbiol 10:691.

12. Sun X, Wang J, Feng D, Ma Z, Li H. 2011. PdCYP51B, a new putative sterol 14α-demethylase gene of Penicillium digitatum involved in resistance to imazalil and other fungicides inhibiting ergosterol synthesis. Appl Microbiol Biotechnol 91:1107–1119.

13. Dunkel N, Liu TT, Barker KS, Homayouni R, Morschhäuser J, Rogers PD. 2008. A Gain-of-Function Mutation in the Transcription Factor Upc2p Causes Upregulation of Ergosterol Biosynthesis Genes and Increased Fluconazole Resistance in a Clinical *Candida albicans* Isolate. Eukaryot Cell 7:1180–1190.

14. Whaley SG, Zhang Q, Caudle KE, Rogers PD. 2018. Relative Contribution of the ABC Transporters Cdr1, Pdh1, and Snq2 to Azole Resistance in Candida glabrata. Antimicrob Agents Chemother 62:e01070–18.

15. Zhang Y, Mao C, Zhai X, Jamieson PA, Zhang C. 2021. Mutation in *cyp51b* and overexpression of *cyp51a* and *cyp51b* confer multiple resistant to DMIS fungicide prochloraz in *Fusarium fujikuroi*. Pest Manag Sci 77:824–833.

16. Vande Zande P, Gautier C, Kawar N, Maufrais C, Metzner K, Wash E, Beach AK, Bracken R, Maciel EI, Pereira De Sá N, Fernandes CM, Solis NV, Del Poeta M, Filler SG, Berman J, Ene IV, Selmecki A. 2024. Step-wise evolution of azole resistance through copy number variation followed by KSR1 loss of heterozygosity in Candida albicans. PLOS Pathog 20:e1012497.

17. Hartmann FE, Vonlanthen T, Singh NK, McDonald MC, Milgate A, Croll D. 2020. The complex genomic basis of rapid convergent adaptation to pesticides across continents in a fungal plant pathogen. Mol Ecol mec.15737.

18. McDonald MC, Renkin M, Spackman M, Orchard B, Croll D, Solomon PS, Milgate A. 2019. Rapid Parallel Evolution of Azole Fungicide Resistance in Australian Populations of the Wheat Pathogen Zymoseptoria tritici. Appl Environ Microbiol 85:e01908–18.

19. Chen Y-C, Lai M-H, Wu C-Y, Lin T-C, Cheng A-H, Yang C-C, Wu H-Y, Chu S-C, Kuo C-C, Wu Y-F, Lin G-C, Tseng M-N, Tsai Y-C, Lin C-C, Chen C-Y, Huang J-W, Lin H-A, Chung C-L. 2016. The Genetic Structure, Virulence, and Fungicide Sensitivity of Fusarium fujikuroi in Taiwan. Phytopathology® 106:624–635.

20. Song Y, Chen X, Sun J, Bai Y, Jin L, Lin Y, Sun Y, Cao H, Chen Y. 2022. *In Vitro* Determination of Sensitivity of *Fusarium fujikuroi* to Fungicide Azoxystrobin and Investigation of Resistance Mechanism. J Agric Food Chem 70:9760–9768.

21. Baek D, Kim HT. 2021. Genetic Mutation Analysis of Botrytis cinerea Resistant to Boscalid. Korean J Pestic Sci 25:246–254.

22. Liu H, Lee G, Sang H. 2024. Exploring SDHI fungicide resistance in *BOTRYTIS CINEREA* through genetic transformation system and ALPHAFOLD model-based molecular docking. Pest Manag Sci 80:5954–5964.

23. Scherm B, Balmas V, Infantino A, Aragona M, Valente MT, Desiderio F, Marcello A, Phanthavong S, Burgess LW, Rau D. 2019. Clonality, spatial structure, and pathogenic variation in Fusarium fujikuroi from rain-fed rice in southern Laos. PLOS ONE 14:e0226556.

24. Veloukas T, Leroch M, Hahn M, Karaoglanidis GS. 2011. Detection and Molecular Characterization of Boscalid-Resistant *Botrytis cinerea* Isolates from Strawberry. Plant Dis 95:1302–1307.

25. Amezrou R, Ducasse A, Compain J, Lapalu N, Pitarch A, Dupont L, Confais J, Goyeau H, Kema GHJ, Croll D, Amselem J, Sanchez-Vallet A, Marcel TC. 2024. Quantitative pathogenicity and host adaptation in a fungal plant pathogen revealed by whole-genome sequencing. Nat Commun 15:1933.

26. Feurtey A, Lorrain C, McDonald MC, Milgate A, Solomon PS, Warren R, Puccetti G, Scalliet G, Torriani SFF, Gout L, Marcel TC, Suffert F, Alassimone J, Lipzen A, Yoshinaga Y, Daum C, Barry K, Grigoriev IV, Goodwin SB, Genissel A, Seidl MF, Stukenbrock EH, Lebrun M-H, Kema GHJ, McDonald BA, Croll D. 2023. A thousand-genome panel retraces the global spread and adaptation of a major fungal crop pathogen. Nat Commun 14:1059.

27. Pereira D, Croll D, Brunner PC, McDonald BA. 2020. Natural selection drives population divergence for local adaptation in a wheat pathogen. Fungal Genet Biol 141:103398.

28. Spanner R, Taliadoros D, Richards J, Rivera-Varas V, Neubauer J, Natwick M, Hamilton O, Vaghefi N, Pethybridge S, Secor GA, Friesen TL, Stukenbrock EH, Bolton MD. 2021. Genome-Wide Association and Selective Sweep Studies Reveal the Complex Genetic Architecture of DMI Fungicide Resistance in *Cercospora beticola*. Genome Biol Evol 13:evab209.

29. Dutta A, McDonald BA, Croll D. 2022. Combined reference-free and multi-reference approaches uncover cryptic variation underlying rapid adaptation in microbial pathogens. preprint. Genomics.

30. Ballu A, Despréaux P, Duplaix C, Dérédec A, Carpentier F, Walker A-S. 2023. Antifungal alternation can be beneficial for durability but at the cost of generalist resistance. Commun Biol 6:180.

31. Noel ZA, Rojas AJ, Jacobs JL, Chilvers MI. 2019. A High-Throughput Microtiter-Based Fungicide Sensitivity Assay for Oomycetes Using *Z* ′-Factor Statistic. Phytopathology® 109:1628–1637.

32. Sabburg R, Gregson A, Urquhart AS, Aitken EAB, Smith L, Thatcher LF, Gardiner DM. 2021. A method for high-throughput image-based antifungal screening. J Microbiol Methods 190:106342.

33. Garrigues S, Kun RS, Peng M, Gruben BS, Benoit Gelber I, Mäkelä M, De Vries RP. 2021. The Cultivation Method Affects the Transcriptomic Response of Aspergillus niger to Growth on Sugar Beet Pulp. Microbiol Spectr 9:e01064–21.

34. Meyer V, Andersen MR, Brakhage AA, Braus GH, Caddick MX, Cairns TC, De Vries RP, Haarmann T, Hansen K, Hertz-Fowler C, Krappmann S, Mortensen UH, Peñalva MA, Ram AFJ, Head RM. 2016. Current challenges of research on filamentous fungi in relation to human welfare and a sustainable bio-economy: a white paper. Fungal Biol Biotechnol 3:6, s40694–016-0024–8.

35. Berkow EL, Lockhart SR, Ostrosky-Zeichner L. 2020. Antifungal Susceptibility Testing: Current Approaches. Clin Microbiol Rev 33:e00069–19.

36. Marcianò D, Toffolatti SL. 2023. Methods for Fungicide Efficacy Screenings: Multiwell Testing Procedures for the Oomycetes Phytophthora infestans and Pythium ultimum. Microorganisms 11:350.

37. McNab E, Rether A, Hsiang T. 2023. Development of a microplate absorbance assay for assessing fungicide sensitivity of filamentous fungi and comparison to an amended agar assay. J Microbiol Methods 204:106653.

38. Fones H, Gurr S. 2015. The impact of Septoria tritici Blotch disease on wheat: An EU perspective. Fungal Genet Biol 79:3–7.

39. Jørgensen LN, Heick TM. 2021. Azole Use in Agriculture, Horticulture, and Wood Preservation – Is It Indispensable? Front Cell Infect Microbiol 11:730297.

40. Torriani SFF, Melichar JPE, Mills C, Pain N, Sierotzki H, Courbot M. 2015. Zymoseptoria tritici: A major threat to wheat production, integrated approaches to control. Fungal Genet Biol 79:8–12.

41. Singh NK, Karisto P, Croll D. 2021. Population-level deep sequencing reveals the interplay of clonal and sexual reproduction in the fungal wheat pathogen Zymoseptoria tritici. Microb Genomics 7:000678.

42. Croll D, Zala M, McDonald BA. 2013. Breakage-fusion-bridge Cycles and Large Insertions Contribute to the Rapid Evolution of Accessory Chromosomes in a Fungal Pathogen. PLoS Genet 9:e1003567.

43. Goodwin SB, Ben M’Barek S, Dhillon B, Wittenberg AHJ, Crane CF, Hane JK, Foster AJ, Van der Lee TAJ, Grimwood J, Aerts A, Antoniw J, Bailey A, Bluhm B, Bowler J, Bristow J, van der Burgt A, Canto-Canché B, Churchill ACL, Conde-Ferràez L, Cools HJ, Coutinho PM, Csukai M, Dehal P, De Wit P, Donzelli B, van de Geest HC, van Ham RCHJ, Hammond-Kosack KE, Henrissat B, Kilian A, Kobayashi AK, Koopmann E, Kourmpetis Y, Kuzniar A, Lindquist E, Lombard V, Maliepaard C, Martins N, Mehrabi R, Nap JPH, Ponomarenko A, Rudd JJ, Salamov A, Schmutz J, Schouten HJ, Shapiro H, Stergiopoulos I, Torriani SFF, Tu H, de Vries RP, Waalwijk C, Ware SB, Wiebenga A, Zwiers L-H, Oliver RP, Grigoriev IV, Kema GHJ. 2011. Finished Genome of the Fungal Wheat Pathogen Mycosphaerella graminicola Reveals Dispensome Structure, Chromosome Plasticity, and Stealth Pathogenesis. PLoS Genet 7:e1002070.

44. Plissonneau C, Stürchler A, Croll D. 2016. The Evolution of Orphan Regions in Genomes of a Fungal Pathogen of Wheat. mBio 7:e01231–16.

45. Croll D, Lendenmann MH, Stewart E, McDonald BA. 2015. The Impact of Recombination Hotspots on Genome Evolution of a Fungal Plant Pathogen. Genetics 201:1213–1228.

46. Stukenbrock EH, Dutheil JY. 2018. Fine-Scale Recombination Maps of Fungal Plant Pathogens Reveal Dynamic Recombination Landscapes and Intragenic Hotspots. Genetics 208:1209–1229.

47. Abraham LN, Oggenfuss U, Croll D. 2024. Population-level transposable element expression dynamics influence trait evolution in a fungal crop pathogen. mBio 15:e02840–23.

48. Oggenfuss U, Badet T, Wicker T, Hartmann FE, Singh NK, Abraham L, Karisto P, Vonlanthen T, Mundt C, McDonald BA, Croll D. 2021. A population-level invasion by transposable elements triggers genome expansion in a fungal pathogen. eLife 10:e69249.

49. Hellin P, Duvivier M, Heick TM, Fraaije BA, Bataille C, Clinckemaillie A, Legrève A, Jørgensen LN, Andersson B, Samils B, Rodemann B, Berg G, Hutton F, Garnault M, El Jarroudi M, Couleaud G, Kildea S. 2021. Spatio-temporal distribution of DMI and SDHI fungicide resistance of *ZYMOSEPTORIA TRITICI* throughout EUROPE based on frequencies of key target-site alterations. Pest Manag Sci 77:5576–5588.

50. Huf A, Rehfus A, Lorenz KH, Bryson R, Voegele RT, Stammler G. 2018. Proposal for a new nomenclature for *CYP51* haplotypes in *Zymoseptoria tritici* and analysis of their distribution in Europe. Plant Pathol 67:1706–1712.

51. Kildea S, Dooley H, Byrne S. 2023. A note on the impact of CYP51 alterations and their combination in the wheat pathogen Zymoseptoria tritici on sensitivity to the azole fungicides epoxiconazole and metconazole. Ir J Agric Food Res 62.

52. Leroux P, Walker A-S. 2011. Multiple mechanisms account for resistance to sterol 14α-demethylation inhibitors in field isolates of Mycosphaerella graminicola. Pest Manag Sci 67:44– 59.

53. Stammler G, Carstensen M, Koch A, Semar M, Strobel D, Schlehuber S. 2008. Frequency of different CYP51-haplotypes of Mycosphaerella graminicola and their impact on epoxiconazole-sensitivity and -field efficacy. Crop Prot 27:1448–1456.

54. Cools HJ, Mullins JGL, Fraaije BA, Parker JE, Kelly DE, Lucas JA, Kelly SL. 2011. Impact of Recently Emerged Sterol 14α-Demethylase (CYP51) Variants of Mycosphaerella graminicola on Azole Fungicide Sensitivity. Appl Environ Microbiol 77:3830–3837.

55. Fraaije BA, Cools HJ, Kim S-H, Motteram J, Clark WS, Lucas JA. 2007. A novel substitution I381V in the sterol 14?-demethylase (CYP51) of Mycosphaerella graminicola is differentially selected by azole fungicides. Mol Plant Pathol 8:245–254.

56. Neau E, Patry-Leclaire S, Duplaix C, Walker A-S, Fillinger S. 2023. Screen of potential multi-drug resistant Zymoseptoria tritici field isolates reveals genotypic and phenotypic diversity suggesting multiple mechanisms involved in MDR16 TH EUROPEAN CONFERENCE ON FUNGAL GENETICS. Susanne Zeilinger-Migsich and Hubertus Haas, Innsbruck,, Austria.

57. Omrane S, Audéon C, Ignace A, Duplaix C, Aouini L, Kema G, Walker A-S, Fillinger S. 2017. Plasticity of the MFS1 Promoter Leads to Multidrug Resistance in the Wheat Pathogen Zymoseptoria tritici. mSphere 2:e00393–17.

58. Eurostat. 2024. Pesticide sales. aei_fm_salpest09.

59. European Commision. 2024. Active substances.

60. De Ramón-Carbonell M, López-Pérez M, González-Candelas L, Sánchez-Torres P. 2019. PdMFS1 Transporter Contributes to Penicilliun digitatum Fungicide Resistance and Fungal Virulence during Citrus Fruit Infection. J Fungi 5:100.

61. Vu BG, Stamnes MA, Li Y, Rogers PD, Moye-Rowley WS. 2021. The Candida glabrata Upc2A transcription factor is a global regulator of antifungal drug resistance pathways. PLOS Genet 17:e1009582.

62. Leroux P, Gredt M, Leroch M, Walker A-S. 2010. Exploring Mechanisms of Resistance to Respiratory Inhibitors in Field Strains of *Botrytis cinerea*, the Causal Agent of Gray Mold. Appl Environ Microbiol 76:6615–6630.

63. Leroux P, Albertini C, Gautier A, Gredt M, Walker A-S. 2007. Mutations in theCYP51 gene correlated with changes in sensitivity to sterol 14α-demethylation inhibitors in field isolates ofMycosphaerella graminicola. Pest Manag Sci 63:688–698.

64. Uffelmann E, Huang QQ, Munung NS, De Vries J, Okada Y, Martin AR, Martin HC, Lappalainen T, Posthuma D. 2021. Genome-wide association studies. Nat Rev Methods Primer 1:59.

65. Bertelsen JR, De Neergaard E, Smedegaard-Petersen V. 2001. Fungicidal effects of azoxystrobin and epoxiconazole on phyllosphere fungi, senescence and yield of winter wheat. Plant Pathol 50:190–205.

66. Schuster M, Kilaru S, Steinberg G. 2024. Azoles activate type I and type II programmed cell death pathways in crop pathogenic fungi. Nat Commun 15:4357.

67. Francisco CS, Ma X, Zwyssig MM, McDonald BA, Palma-Guerrero J. 2019. Morphological changes in response to environmental stresses in the fungal plant pathogen Zymoseptoria tritici. Sci Rep 9:9642.

68. Zwiers L-H, Stergiopoulos I, Van Nistelrooy JGM, De Waard MA. 2002. ABC Transporters and Azole Susceptibility in Laboratory Strains of the Wheat Pathogen *Mycosphaerella graminicola*. Antimicrob Agents Chemother 46:3900–3906.

69. Copier C, Osorio-Navarro C, Maldonado JE, Auger J, Silva H, Esterio M. 2024. A Conservative Mutant Version of the Mrr1 Transcription Factor Correlates with Reduced Sensitivity to Fludioxonil in Botrytis cinerea. Pathog Basel Switz 13:374.

70. Kretschmer M, Leroch M, Mosbach A, Walker A-S, Fillinger S, Mernke D, Schoonbeek H-J, Pradier J-M, Leroux P, De Waard MA, Hahn M. 2009. Fungicide-Driven Evolution and Molecular Basis of Multidrug Resistance in Field Populations of the Grey Mould Fungus Botrytis cinerea. PLoS Pathog 5:e1000696.

71. Olivares-Yañez C, Sánchez E, Pérez-Lara G, Seguel A, Camejo PY, Larrondo LF, Vidal EA, Canessa P. 2021. A comprehensive transcription factor and DNA-binding motif resource for the construction of gene regulatory networks in Botrytis cinerea and Trichoderma atroviride. Comput Struct Biotechnol J 19:6212–6228.

72. Nishida N, Jing D, Kuroda K, Ueda M. 2014. Activation of signaling pathways related to cell wall integrity and multidrug resistance by organic solvent in Saccharomyces cerevisiae. Curr Genet 60:149–162.

73. Hu M, Chen S. 2021. Non-Target Site Mechanisms of Fungicide Resistance in Crop Pathogens: A Review. Microorganisms 9:502.

74. Hossam Abdelmonem B, Abdelaal NM, Anwer EKE, Rashwan AA, Hussein MA, Ahmed YF, Khashana R, Hanna MM, Abdelnaser A. 2024. Decoding the Role of CYP450 Enzymes in Metabolism and Disease: A Comprehensive Review. Biomedicines 12:1467.

75. Mirzadi Gohari A, Mehrabi R, Robert O, Ince IA, Boeren S, Schuster M, Steinberg G, de Wit PJGM, Kema GHJ. 2014. Molecular characterization and functional analyses of *ZtWor1*, a transcriptional regulator of the fungal wheat pathogen *Zymoseptoria tritici*: Functionality of *ZtWor1*. Mol Plant Pathol 15:394–405.

76. Bolger AM, Lohse M, Usadel B. 2014. Trimmomatic: a flexible trimmer for Illumina sequence data. Bioinformatics 30:2114–2120.

77. Langmead B, Salzberg SL. 2012. Fast gapped-read alignment with Bowtie 2. Nat Methods 9:357–359.

78. McKenna A, Hanna M, Banks E, Sivachenko A, Cibulskis K, Kernytsky A, Garimella K, Altshuler D, Gabriel S, Daly M, DePristo MA. 2010. The Genome Analysis Toolkit: A MapReduce framework for analyzing next-generation DNA sequencing data. Genome Res 20:1297–1303.

79. Danecek P, Auton A, Abecasis G, Albers CA, Banks E, DePristo MA, Handsaker RE, Lunter G, Marth GT, Sherry ST, McVean G, Durbin R, 1000 Genomes Project Analysis Group. 2011. The variant call format and VCFtools. Bioinformatics 27:2156–2158.

80. Dray S, Dufour A-B. 2007. The **ade4** Package: Implementing the Duality Diagram for Ecologists. J Stat Softw 22.

81. Barker BS, Cocio JE, Anderson SR, Braasch JE, Cang FA, Gillette HD, Dlugosch KM. 2019. Potential limits to the benefits of admixture during biological invasion. Mol Ecol 28:100–113.

82. Francis RM. 2017. POPHELPER : an R package and web app to analyse and visualize population structure. Mol Ecol Resour 17:27–32.

83. Purcell S, Neale B, Todd-Brown K, Thomas L, Ferreira MAR, Bender D, Maller J, Sklar P, de Bakker PIW, Daly MJ, Sham PC. 2007. PLINK: A Tool Set for Whole-Genome Association and Population-Based Linkage Analyses. Am J Hum Genet 81:559–575.

84. Hijmans RJ. 2010. geosphere: Spherical Trigonometry.

85. Ingvarsson PK. 2005. Nucleotide Polymorphism and Linkage Disequilibrium Within and Among Natural Populations of European Aspen (Populus tremula L., Salicaceae). Genetics 169:945–953.

86. Remington DL, Thornsberry JM, Matsuoka Y, Wilson LM, Whitt SR, Doebley J, Kresovich S, Goodman MM, Buckler ES. 2001. Structure of linkage disequilibrium and phenotypic associations in the maize genome. Proc Natl Acad Sci 98:11479–11484.

87. Zhou X, Stephens M. 2012. Genome-wide efficient mixed-model analysis for association studies. Nat Genet 44:821–824.

88. Cingolani P, Platts A, Wang LL, Coon M, Nguyen T, Wang L, Land SJ, Lu X, Ruden DM. 2012. A program for annotating and predicting the effects of single nucleotide polymorphisms, SnpEff: SNPs in the genome of Drosophila melanogaster strain w 1118 ; iso-2; iso-3. Fly (Austin) 6:80–92.

89. Voichek Y, Weigel D. 2020. Identifying genetic variants underlying phenotypic variation in plants without complete genomes. Nat Genet 52:534–540.

90. Quinlan AR, Hall IM. 2010. BEDTools: a flexible suite of utilities for comparing genomic features. Bioinformatics 26:841–842.

91. Huson DH, Bryant D. 2006. Application of Phylogenetic Networks in Evolutionary Studies. Mol Biol Evol 23:254–267.

92. Badet T, Oggenfuss U, Abraham L, McDonald BA, Croll D. 2020. A 19-isolate reference-quality global pangenome for the fungal wheat pathogen Zymoseptoria tritici. BMC Biol 18:12.

93. O’Donnell S, Fischer G. 2020. MUM&Co: accurate detection of all SV types through whole-genome alignment. Bioinformatics 36:3242–3243.

94. Li H. 2018. Minimap2: pairwise alignment for nucleotide sequences. Bioinformatics 34:3094– 3100.

95. Kirsche M, Prabhu G, Sherman R, Ni B, Battle A, Aganezov S, Schatz MC. 2023. Jasmine and Iris: population-scale structural variant comparison and analysis. Nat Methods 20:408–417.

96. Garrison E, Sirén J, Novak AM, Hickey G, Eizenga JM, Dawson ET, Jones W, Garg S, Markello C, Lin MF, Paten B, Durbin R. 2018. Variation graph toolkit improves read mapping by representing genetic variation in the reference. Nat Biotechnol 36:875–879.

97. Sirén J, Monlong J, Chang X, Novak AM, Eizenga JM, Markello C, Sibbesen JA, Hickey G, Chang P-C, Carroll A, Gupta N, Gabriel S, Blackwell TW, Ratan A, Taylor KD, Rich SS, Rotter JI, Haussler D, Garrison E, Paten B. 2021. Pangenomics enables genotyping of known structural variants in 5202 diverse genomes. Science 374:abg8871.

98. Auwera G van der, O’Connor BD. 2020. Genomics in the cloud: using Docker, GATK, and WDL in TerraFirst edition. O’Reilly Media, Sebastopol, CA.

99. Grandaubert J, Bhattacharyya A, Stukenbrock EH. 2015. RNA-seq-Based Gene Annotation and Comparative Genomics of Four Fungal Grass Pathogens in the Genus *Zymoseptoria* Identify Novel Orphan Genes and Species-Specific Invasions of Transposable Elements. G3 GenesGenomesGenetics 5:1323–1333.

100. Jones P, Binns D, Chang H-Y, Fraser M, Li W, McAnulla C, McWilliam H, Maslen J, Mitchell A, Nuka G, Pesseat S, Quinn AF, Sangrador-Vegas A, Scheremetjew M, Yong S-Y, Lopez R, Hunter S. 2014. InterProScan 5: genome-scale protein function classification. Bioinforma Oxf Engl 30:1236–1240.

101. Falcon S, Gentleman R. 2007. Using GOstats to test gene lists for GO term association. Bioinformatics 23:257–258.

102. Bowler J, Scott E, Tailor R, Scalliet G, Ray J, Csukai M. 2010. New capabilities for *Mycosphaerella graminicola* research. Mol Plant Pathol 11:691–704.

103. Zarnack K, Maurer S, Kaffarnik F, Ladendorf O, Brachmann A, Kämper J, Feldbrügge M. 2006. Tetracycline-regulated gene expression in the pathogen Ustilago maydis. Fungal Genet Biol 43:727–738.

