## Supplementary Information for "A large European diversity panel reveals complex azole fungicide resistance gains of a major wheat pathogen"

### Supplementary Figures

### Supplementary Tables

### Supplementary Data

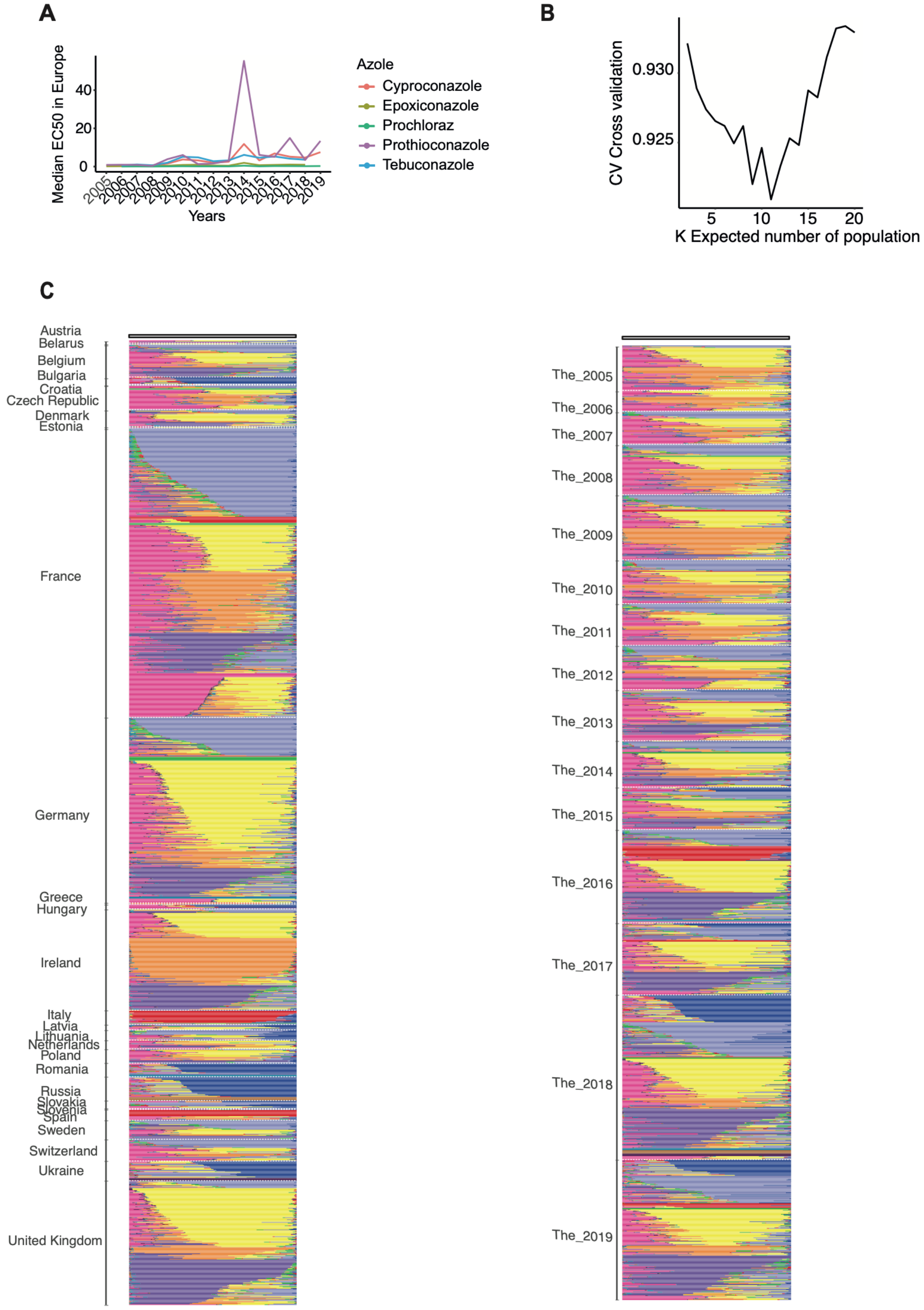


**Supplementary Figure S1**. (A) Overview of the variation in fungicide resistance over time at the European scale. Median $EC_{50}$ values were calculated for five distinct DMI across Europe for each year from 2005 to 2019. (B) Optimal number of genetic clusters (admixture groups), determined by evaluating cross-validation (CV) scores. The lowest CV value indicates the best-fitting admixture model. (C–D) Detailed results from the admixture assignments based on 11 clusters.

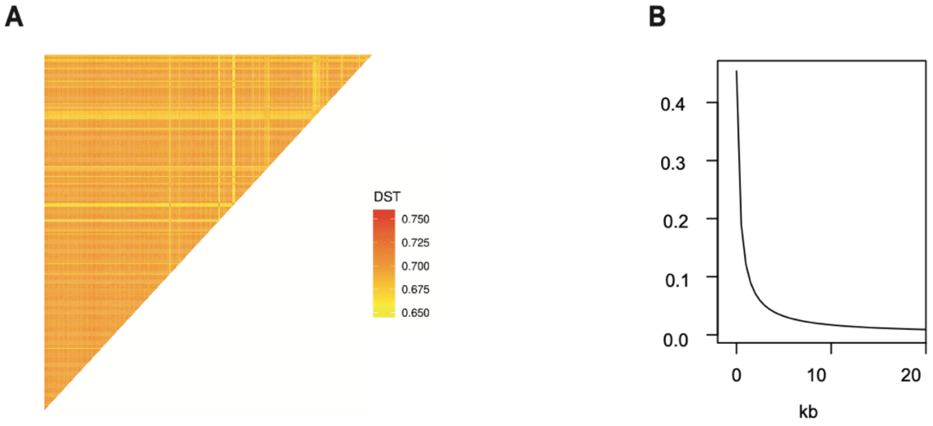


**Supplementary Figure S2**. (A) Heatmap of the identity-by-state (IBS) distances in the European diversity panel. Linkage disequilibrium decay expressed as $r^2$ is shown over physical distances for chromosome 1.

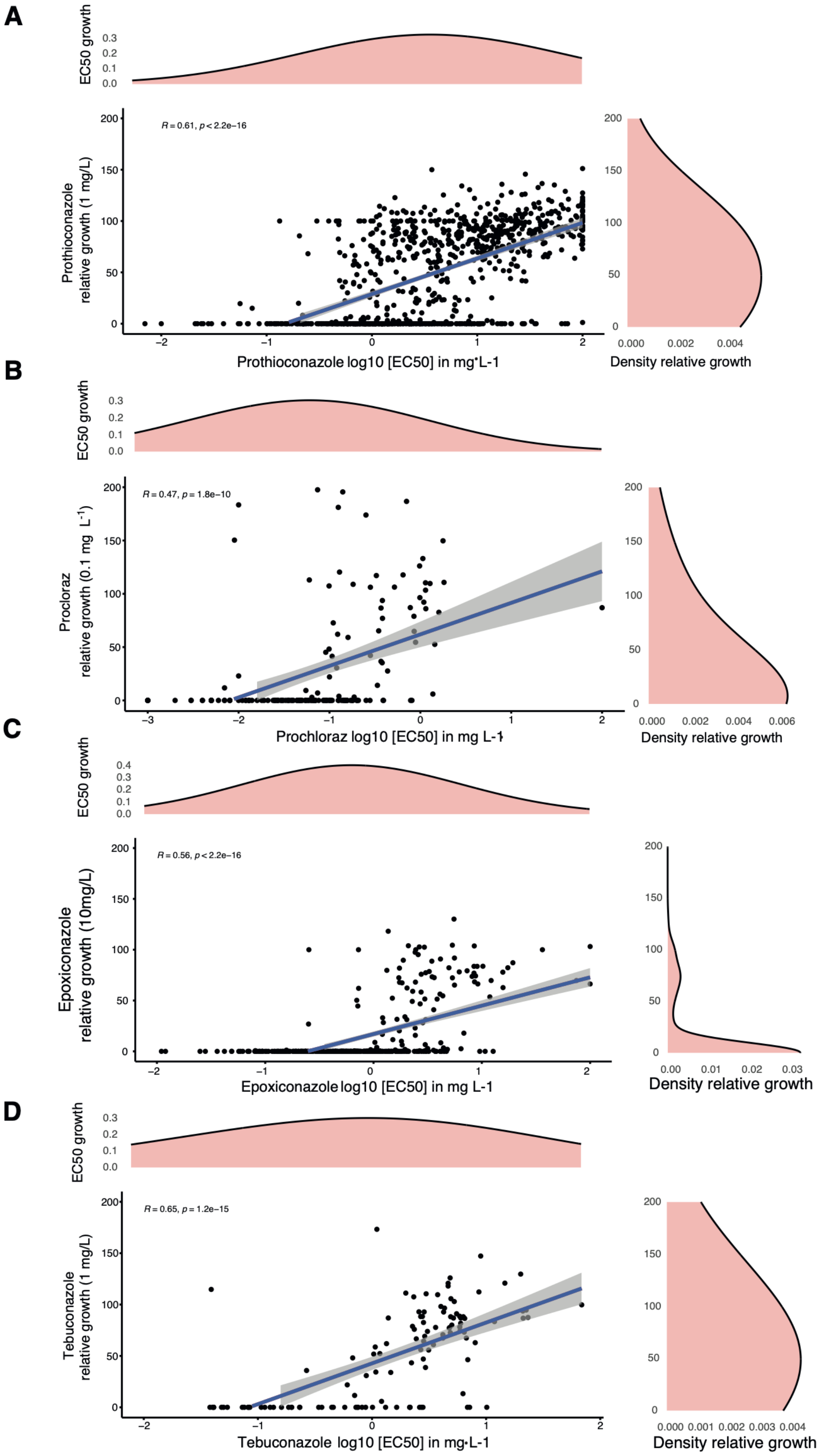


**Supplementary Figure S3**. Relationship between $EC_{50}$ values and relative growth under (A) prothioconazole, (B) prochloraz, (C) epoxiconazole and (D) tebuconazole treatment. A linear model following the formula log10(x) for $EC_{50}$ values was used to fit the trend line. Densities of $EC_{50}$ and relative growth values are shown.

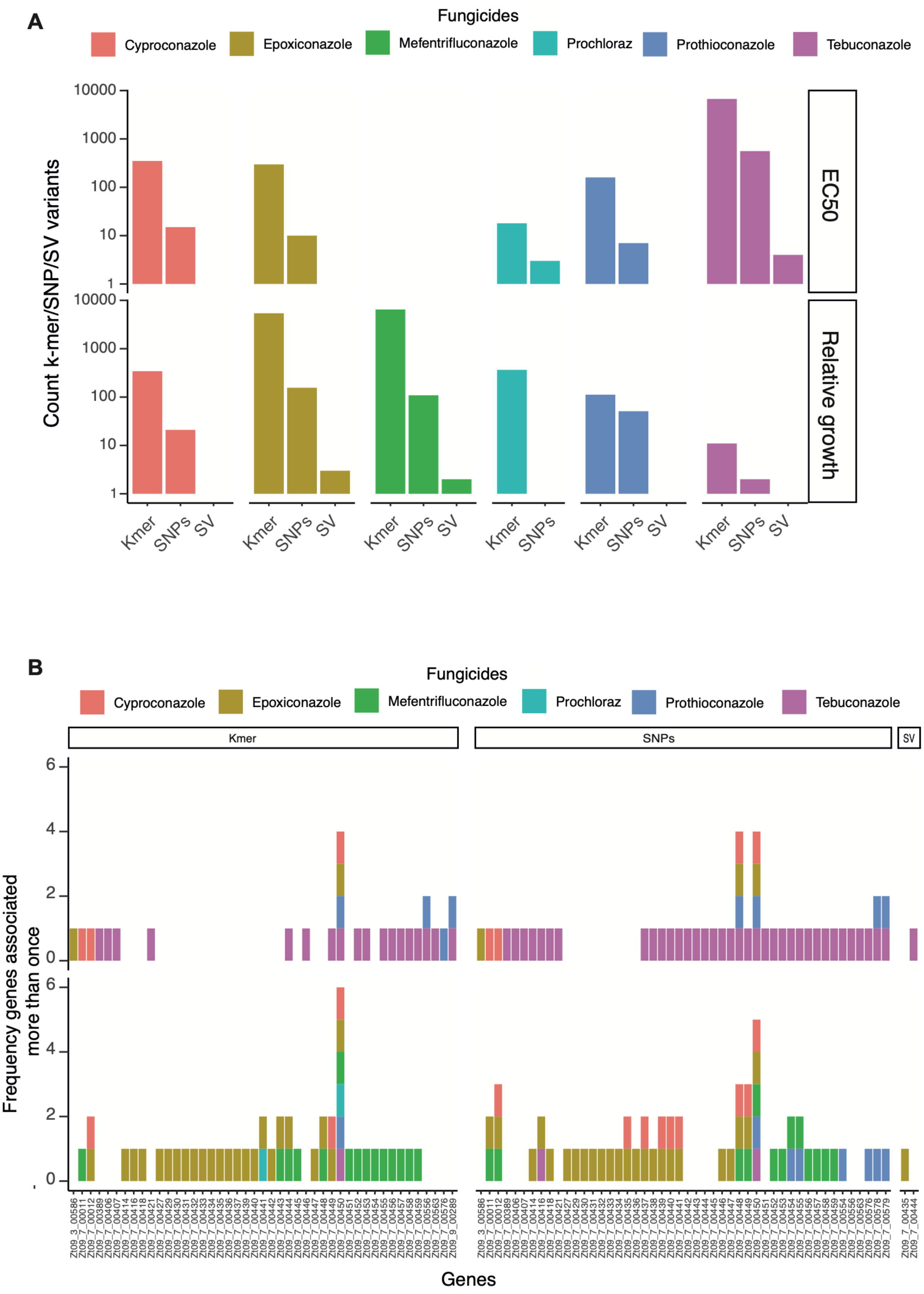


**Supplementary Figure S4**. (A) Total number of significant k-mers, SNPs and SVs identified across the 6 tested DMIs. (B) Genes associated to DMIs resistance across multiple analyses: fungicides (6 DMIs), GWAS approach (SVs, k-mers, SVs), resistance assays ($EC_{50}$, relative growth).

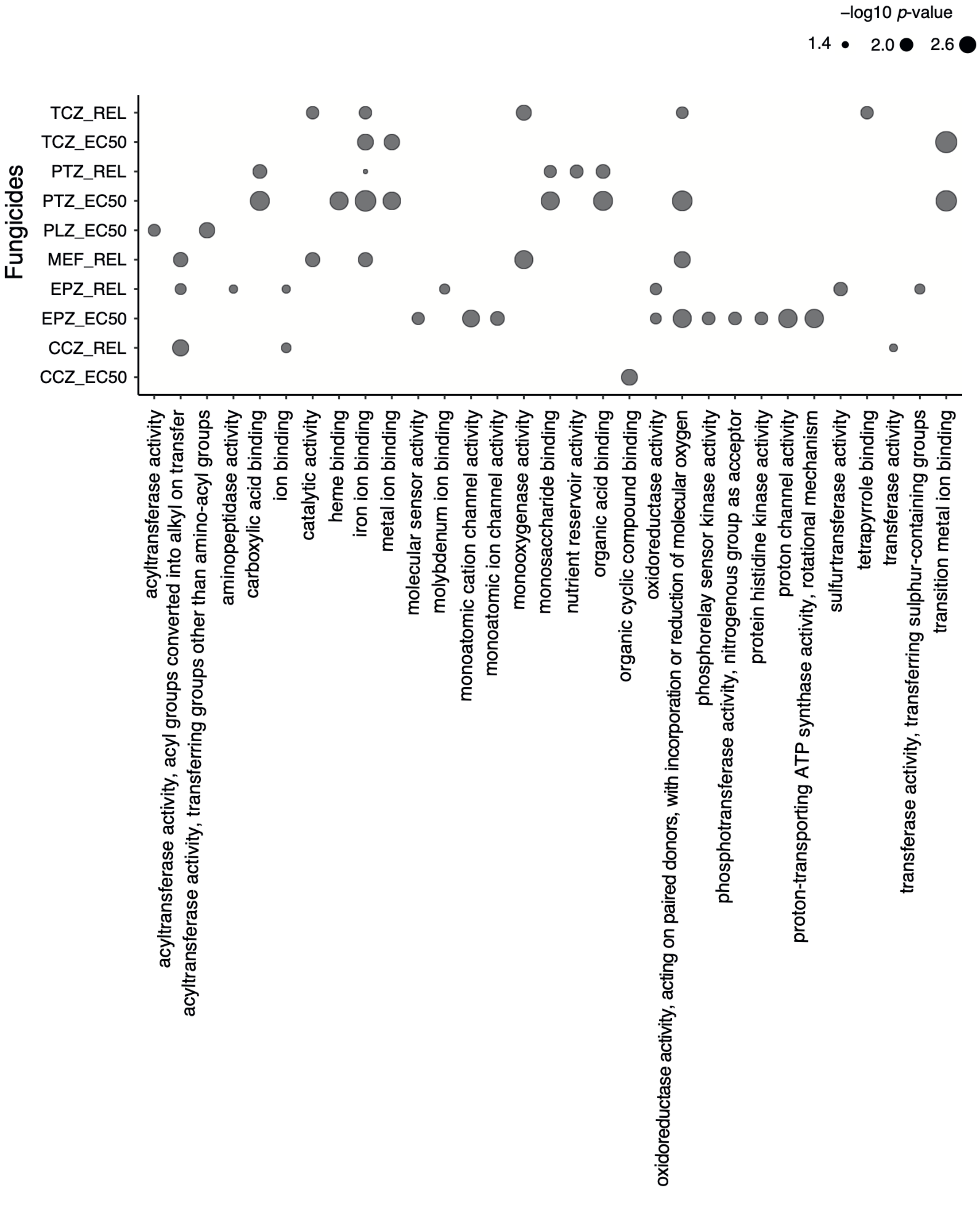


**Supplementary Figure S5**: (A) Enrichment analyses *p*-values of gene functions associated through GWAS for $EC_{50}$ and relative growth values per fungicide.

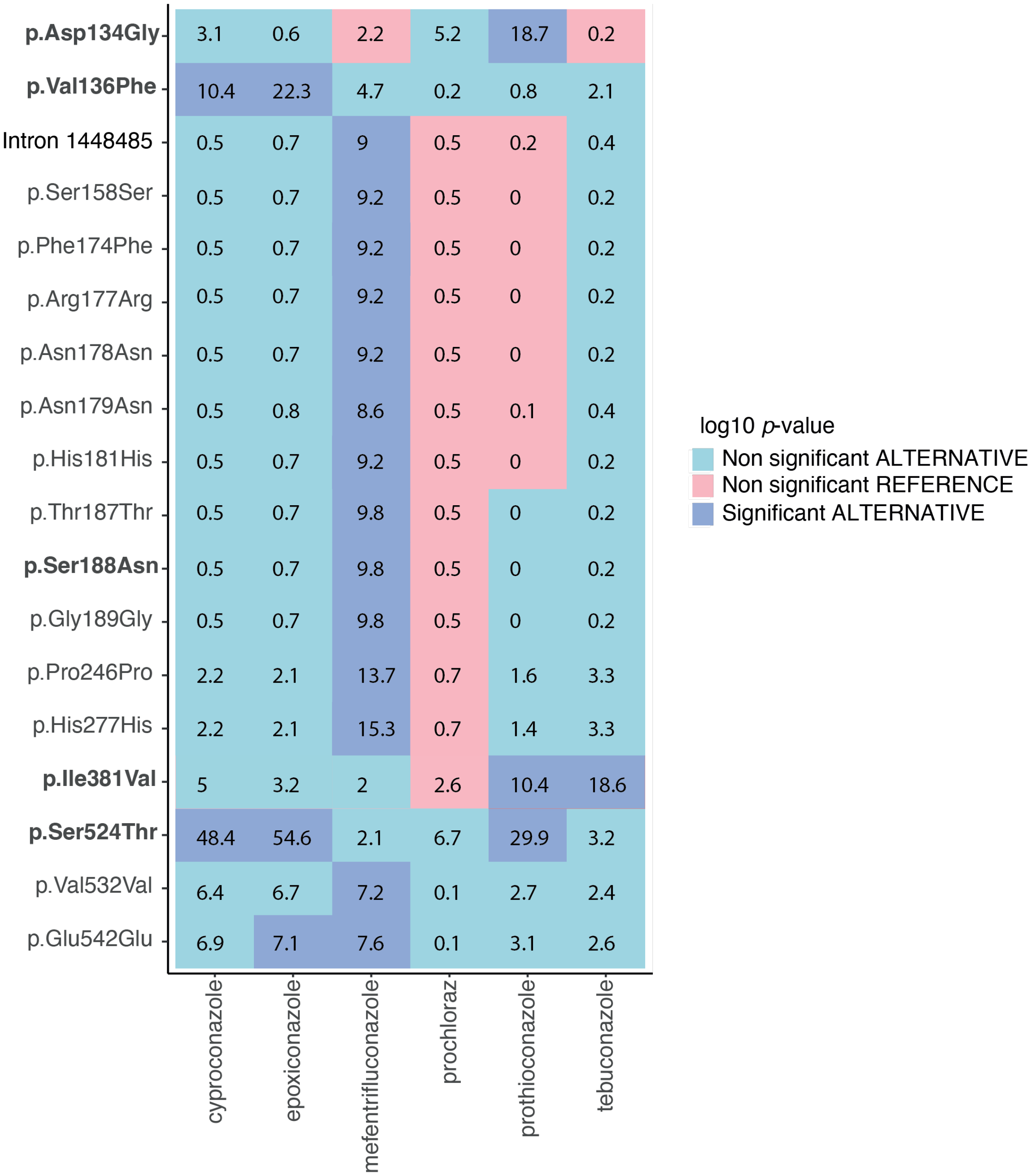


**Supplementary Figure S6**: Heatmap of SNPs significantly associated to DMI resistance based on relative growth values. In blue, the significant alternative alleles (*i.e.* distinct form the IPO323 reference genome representing a susceptible isolate), in light blue and pink the non-significantly associated reference and alternative alleles.

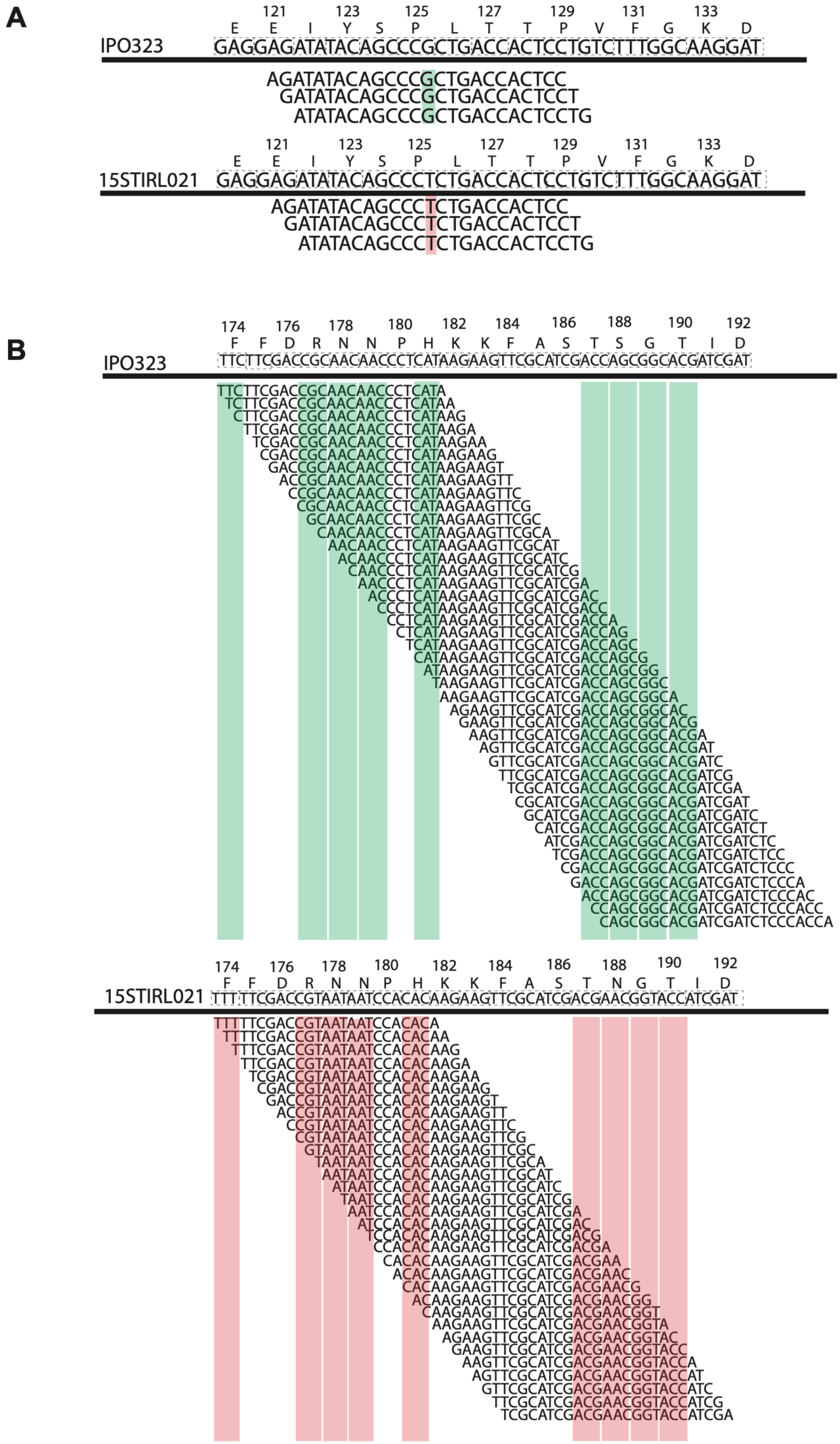

**Supplementary Figure S7**: K-mers significantly associated to mefentrifluconazole and tebuconazole $EC_{50}$ values aligned to the reference genome (IPO323) in the region stretching from 1,448,572-1,448,574 bp (Zone 3) which includes the Pro125Pro synonymous substitution of the *Cyp51* gene. In green, the k-mers positively associated to resistance, in red positively associated to susceptibility. (B) K-mers significantly associated to mefentrifluconazole and tebuconazole $EC_{50}$ from 1,448,318-1,448,359 bp overlapping the codons Phe174, Arg177, Asn178, Asn179, His181, Thr187, Ser188, Gly189 (zone 4) of *Cyp51*. In green, variants positively associated to susceptibility, in red, positively associated to resistance.

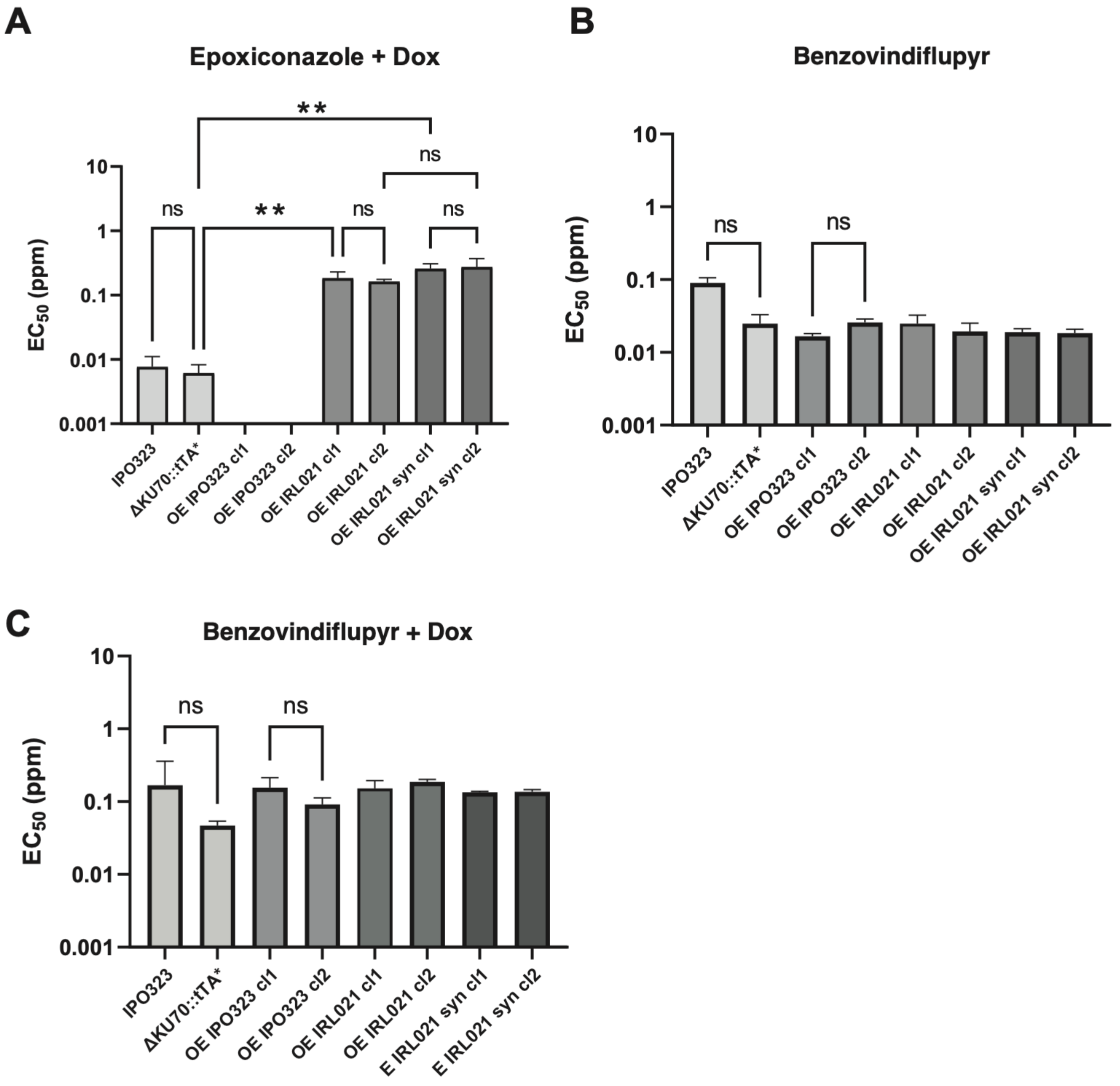


**Supplementary Figure S8**: $EC_{50}$ values of the strains/mutants IPO323 (wild-type), ΔKU70::tTA, OE_IPO323, OE_IRL21 and OE_IRL21SYN in liquid YPD media amended with (A) epoxiconazole and supplemented with doxycycline; (B) benzovindiflupyr and (C) benzovindiflupyr supplemented with doxycycline. One-way ANOVA: **** *p*-value<0.0001, ** *p*-value<0.05.

### Supplementary Table legends

(Available from Zenodo: https://doi.org/10.5281/zenodo.15052606)

**Table S1**: Overview of all *Z. tritici* isolates analyzed in this study. The columns reporting phenotyping information consist of EC50 and relative growth values. EC50 values are the column with 'EC50' followed by an abbreviation for the fungicide: CCZ for cyproconazole, EPZ for epoxiconazole, PLZ for prochloraz, PTZ for prothioconazole, and TCZ for tebuconazole. For relative growth values, the column consists of a combination of acronyms: "Rel_" standing for "relative growth" followed by the same fungicide abbreviation, the fungicide concentration and the day at which the measurement was done (7dpi). The column "Contributing_members_names" lists the names of the contributing members or authors who provided data or were involved in the generation of the isolates. "Used_in_PCA" lists all the strains used to produced the overall PCA of the global pathogen diversity. Finally "Strains_used_in_kmer_GWAS" and "Strains_used_in_SV_GWAS" consist of the strains included in the k-mer and SV analysis, respectively.

**Table S2**: Summary of fungicide resistance assessed by GWAS with corresponding association mapping output files listed. The column "Fungicide_ID_in_GWAS_studies" shows the unique ID shown in Table S1. The column "SNPs_GWAS_tables", "SV_GWAS_tables" and "Pass_threshold_5_Kmers_GWAS" correspond to GEMMA output of the association mapping and available as Supplementary Data 3 from Zenodo (https://doi.org/10.5281/zenodo.15052606).

**Table S3**: Results of association mapping for significant SNPs, k-mers and SVs crossing the Bonferroni threshold across 11 tested traits. The column "beta" and "p_wald" correspond to GEMMA software output (Zhou and Stephens 2012). The column "gene" provides information of the gene intersecting SV, k-mer and SNP variants. Fungicide_ID_in_GWAS_studies corresponds to the fungicide ID (Table S1). The column "mismatch" is specific for k-mers GWAS associations and corresponds to the position and allelic variation obtained from kmer mapping. The column "Kmer_for_match" corresponds to the k-mer sequences on the positive strand, while "Kmer_output" corresponds to the k-mer output name of the k-mer GWAS, and "strand" specifies the DNA strand (forward or reverse compared to the reference genome sequence) where the k-mer is located. The column "Genotyping_approach" refers to the genotyping method used to identify genetic variants. "HGVS.p" is specific for SNP variants and denotes the variant annotation in HGVS format, describing the change in protein sequence obtained through SnpEff. The "Annotation REF" column provides information regarding the reference allele for SVs and SNPs based on the IPO323 genome annotation.

**Table S4**: provides detailed information on SNPs and indels identified within the *Cyp51* coding sequence. The "Alternative" column identifies the alternative allele for each SNP. The "HGVS.p" column provides a description of the amino acid variants. The "Variants" column describes the amino acid variant type. "Variants_kept_for_analysis" indicates SNPs were retained for GWAS.

**Table S5**: Haplotype profile of the *Cyp51* gene based on significantly associated alleles and their frequencies. The 'Haplotype' and "Haplotype number" columns represent the combination of missense and synonymous amino acid changes, indicating whether a combination was associated with resistance (1), susceptibility (0), or not associated (NA) based on the GWAS. The 'frequency_haplotypes' column reports haplotype frequencies in the European diversity panel. The 'Resistant_count' column indicates the total number of resistant alleles per haplotype.

**Table S6**: Significance values for 17 associated *Cyp51* SNPs for fungicide resistance assessed by relative growth. The predicted impact of each allele as identified by SnpEff is shown with "HGVS.p" showing the HGVS notation. "Putative_impact" is an assessment of the impact or deleteriousness of the SNP, categorized as {HIGH, MODERATE, LOW, MODIFIER}. The column "Annotation" describes the gene region where the SNP was identified. The column "isolate" corresponds to the isolate name and “GT” the genotype (1 alternative, 0 reference SNP allele), finally the column “Resistant_susceptible” reports whether the strain is resistant or susceptible according to GWAS.

**Table S7**: EC50 assessments of the isolate IPO323, and mutants ΔKU70::tTA*, OE IPO323 cl1, OE IPO323 cl2, OE IRL021 cl1, OE IRL021 cl2, OE IRL021SYN cl1 and OE IRL021SYN cl2 in mefentrifluconazole, epoxiconazole and benzovindiflupyr. Each of the EC50 values represent the mean value of three technical replicates. "Nd" indicates not determined EC50 values.

**Table S8**: Different culture media used in this study.

**Table S9**: Overview of the assayed fungicides. "Study's purpose" outlines the study objectives for each fungicide such as GWAS or functional validation. The column “media” details the experimental setups used, including solid agar plates or liquid culture systems. The “Fungicide resistance measurement” details how resistance levels were quantified. The “Concentration**”** and **“**dpi**”** specific the concentrations and days post inoculation at which measurements were made. The column “Fungicide_ID_in_GWAS_studies” correspond to the unique IDs shown in Table S1. The columns “Pesticide use in EU (in the time frame 2005–2019)” reports if the fungicide applications across EU countries covered the specified timeframe. The "Pesticide used in wheat in EU" identifies whether the fungicides were specifically applied to wheat crops during the study period. “Reported resistance *Zymoseptoria tritici*" reports whether resistance to the specified fungicides had been reported prior to this study. Citation columns provide key references.

**Table S10**: Pairwise genetic distances and geographic locations of European diversity panel isolates. The columns “IID1” and “IID2”**:** correspond to the pair of isolates analyzed. The column “DST” reports the genetic relatedness between the two individuals. “Latitude_IID1” and “Longitude_IID1” correspond to latitude and longitude of the first isolate. “Latitude_IID2” and “Longitude_IID2” correspond to latitude and longitude of the second isolate. Finally, “distance”, correspond to the geographic distance (in m) between the two locations.

**Table S11**: Proportion of variance explained (PVE) by genotyping methods and different fungicides analyzed based on GEMMA. Each row represents the results for a specific fungicide, detailing the PVE, proportion of variance explained, the standard error of the PVE (pve_se), the genotyping method and the fungicide used.

**Table S12**: Correlation analyses of relative growth and EC50 values used for fungicide resistance assessments. The columns “Fungicide_1” and “Fungicide_2” identifies the pairs of fungicides analyzed for correlations. Pearson's correlation coefficient and sample size, Student's $t$-statistic and $p$-value.

**Table S13**: Three haplotypes of the *CYP51* sequence for transformations. The column “Name mutant” and “Name plasmid”, refers to the names of the strains and plasmid vector used to in this study. Finally the column “gb_file” refers to the GenBank file and contains annotations of the gene locations, function, sequence information, about the plasmid and inserted sequence.

**Table S14**: Primers used to verify homologous recombination events of OE_IRL21SYN, OE_IRL21 and OE_IPO323 inserts into the IPO323 background. "Position" indicates the matching of the oligo annealing on the target sequence while "OLIGO" corresponds to the name of the primer. "Len" is the length of the oligonucleotide (in bp) and "Tm" is the melting temperature (°C) of the oligo. "GC%" is reported for the oligo. The column "Any_th", refers to potential binding of the oligo to non-target sequences. The column "3_th", describe the specificity at the 3' end of the oligo. The column "Hairpin" indicates the presence or absence of hairpin structures within the oligo. Finally "Seq" provides the nucleotide sequence of the oligonucleotide. Further information include the PCR polymerase used either with Q5 (Q5® High-Fidelity DNA Polymerase - New England Biolabs) or G2 (GoTaq® G2 Flexi DNA Polymerase - Promega).

### Supplementary datasets

(Available from Zenodo: https://doi.org/10.5281/zenodo.15052606)

**Supplementary Data 1**: Plasmid_Ku70: The Plasmid_Ku70 construct includes the pNOV2114 backbone fused with Ku70, a TetRep (tetracycline repressor), an NPT2 (neomycin phosphotransferase) resistance gene.

**Supplementary Data 2**: Plasmids.gb: include AltsdhC_Hygr_pNOV2114_CYP51_IPO32, AltsdhCHygr_pNOV2114_CYP51_IRL021 and AltsdhC_Hygr_pNOV2114_CYP51_IRL021_SYN. The construct contains the different *cyp51* haplotypes used in the functional validation. The hygromycin resistance (Hygr) marker is present to allow for selection in culture. The *pNOV2114* plasmid serves as the backbone vector for delivering this genetic modification.

**Supplementary Data 3:** Association mapping data files detailed in Supplementary Table S2.
- SNPs_* tables: Repository of all SNPs association tables generated with GEMMA.
- SV_* tables: Repository of all SV association tables generated with GEMMA.
- Kmer_* tables: Repository of significant K-mer associations.
